# Mutant p53 Directs PARP to Regulate Replication Stress and Drive Breast Cancer Metastasis

**DOI:** 10.64898/2026.03.26.713220

**Authors:** Gu Xiao, George K. Annor, Katherine W. Harmon, Valery Chavez, Fayola Levine, Samuel Ahuno, Mohamed I. Atmane, Samantha St. Jean, Florencia P. Madorsky Rowdo, Pamella Leybengrub, AnnMarie Gaglio, Viola Ellison, Divya Venkatesh, Siyu Sun, Taha Merghoub, Benjamin Greenbaum, Olivier Elemento, Melissa B. Davis, Olorunseun Ogunwobi, Jill Bargonetti

## Abstract

*TP53* mutations occur in 80-90% of triple-negative breast cancers (TNBCs) and drive genomic instability and metastatic progression. Poly (ADP-ribose) polymerase (PARP) is critical for DNA repair and replication fork stability. How oncogenic signaling influences PARP function to sustain proliferation during replication stress remains unclear. Mutant p53 (mtp53) R273H associates tightly with chromatin, forms complexes with PARP, and enhances PARP recruitment to replication forks [1–3]. The C-terminal region of mtp53 mediates mtp53-PARP and mtp53-Poly (ADP-ribose) (PAR) interactions that facilitate S phase progression [4, 5]. The PARP inhibitor talazoparib (TAL) combined with the alkylating agent temozolomide (TMZ) produces synergistic cytotoxicity selectively in mtp53, but not wild-type p53 (wtp53), breast cancer cells and organoids. Herein we evaluated the mechanism of mtp53-associated cell death and tested if this could translate to a preclinical xenograft model. We found that TMZ+TAL treatment induced elevated cleaved PARP and γH2AX and reduced the metastasis-promoting oncoprotein MDMX. In orthotopic xenografts expressing mtp53 R273H, but not wtp53, combination therapy significantly decreased circulating tumor cells (CTCs) and lung metastases. Transcriptomic profiling of tumors from combination treated animals demonstrated downregulation of *MDMX*, *VEGF,* and *NF-κB*, consistent with the observed suppression of CTCs and lung metastasis, and increased γH2AX, indicative of replication stress in mtp53 xenografts. Inhibition of metastasis was also observed in mtp53 R273H WHIM25 and p53-undetectable WHIM6 TNBC patient-derived xenografts (PDX). The mtp53 C-terminal domain (347-393) demonstrated a critical tumor promoting function, as CRISPR-mediated deletion impaired replication fork progression, tumor growth, and metastatic dissemination. DNA fiber combing showed that expression of full-length mtp53 R273H, but not C-terminal deleted Δ347-393, supported sustained single-stranded DNA gaps (ssGAPs) following Poly (ADP-ribose) glycohydrolase (PARG) inhibition. These findings support that mtp53 uses C-terminal amino acids to exploit PARP to enable replication stress adaptation and that mtp53 is a predictive biomarker for combined PARP inhibitor and DNA damaging therapies targeting TNBC.

**Significance statement:** *TP53* mutations are the most common genetic alterations in TNBC and a major driver of replication stress and metastasis. This study shows that missense mutant p53 uses C-terminal amino acids to reprogram PARP activity to maintain tumor cell survival under replication stress. We demonstrate that p53 status governs the response to combined PARP inhibitor (PARPi) and DNA-damaging chemotherapy, establishing an additional molecular basis beyond BRCA1 mutations for treating TNBC with PARPi therapy. These findings reveal a previously unrecognized mechanism by which the mutant p53-PARP axis enables replication stress tolerance and drives cancer metastasis. We show mutation of p53 in TNBC provides an additional biomarker-guided framework to improve PARPi therapeutic outcomes.

## Introduction

*TP53* mutations occur in approximately 80-90% of triple-negative breast cancers (TNBCs) [6], and mutant p53 (mtp53) is a major driver of genomic instability and metastatic progression [7, 8]. Cancer cells frequently encounter obstacles to replication fork progression and defects in the DNA damage response (DDR), resulting in incomplete DNA repair, aberrant replication, and elevated replication stress [9]. Replication stress arises when fork progression is impeded by DNA damage, difficult-to-replicate regions, transcription-replication conflicts, nucleotide shortage, or oncogene activation [10]. Replication stress can cause fork stalling, fork reversal, accumulation of single-stranded DNA (ssDNA), and genomic instability [11]. Unlike wild-type p53, which activates genes to arrest the cell cycle or stimulate repair in response to stress, mtp53 acquires oncogenic gain-of-function (GOF) activities that promote tumorigenesis through both transcription-dependent and -independent mechanisms [7, 12]. These GOF functions enable tumor cells with mtp53 to tolerate chronic replication stress by reprogramming DNA repair and replication pathways, thereby sustaining proliferation under genotoxic conditions [13]. Notably, mtp53 perturbs replication checkpoint control by enhancing the TopBP1-Treslin interaction, leading to unscheduled origin firing and fork instability, while upregulating CDC7 activity to drive hyper-origin firing, further exacerbating replication stress [14, 15]. Thus, mtp53 amplifies replication stress while reconfiguring DNA damage tolerance pathways to maintain continuous DNA synthesis.

Our previous studies revealed that the hot spot missense mutation in p53 at amino acid (AA) position 273 (R273H) produces stable protein that is chromatin-associated and in close proximity with replicative helicase Minichromosome Maintenance (MCM) complex, Poly(ADP-ribose) polymerase PARP and poly(ADP-ribose) (PAR), promoting PARP recruitment to replication forks [1–3, 5]. Analysis of The Cancer Genome Atlas (TCGA) shows significant co-elevation of p53 and PARP protein levels in TNBC [3]. It has been shown that at replication forks, PARP coordinates with replication stress response (RSR), stabilizing stalled forks through MRE11 recruitment and regulating fork restart by suppressing RECQ1 helicase [16–19]. In addition, PARP recruits XRCC1 and DNA ligase III (LIG3), critical for Okazaki fragment maturation [20]. Consistent with these functions, S-phase represents the principal period for PARylation in proliferating cells, with poly(ADP-ribose) glycohydrolase (PARG) degrading PAR to prevent toxic accumulation of PAR polymers [21]. Furthermore, we demonstrated that the C-terminal domain (CTD) of mtp53 directly binds PARP and PAR, orchestrating PARP-dependent replication stress tolerance and sustaining DNA synthesis [4, 5]. Deletion of the mtp53 CTD impairs TNBC cell proliferation and S-phase progression, highlighting the role of specific regions of mtp53 involved in replication stress adaptation [4, 5].

MDMX (MDM4) and MDM2 are structurally related regulators of wtp53. MDMX enhances MDM2 E3 ligase activity to ubiquitinates wtp53 for degradation [22]. In addition, MDMX and MDM2 contribute to metastasis by activating CXCR4, facilitating circulating tumor cell dissemination and lung metastasis in mtp53 breast cancer cells [23]. Furthermore, MDMX establishes a positive feedback loop in which MDMX and the CXCL12-CXCR4 axis reinforce one another to promote metastasis [24]. Additionally, mtp53 drives metastasis through epithelial-to-mesenchymal transition (EMT) by activating NF-κB, TGF-β, and integrin/FAK signaling, and upregulating transcription factors such as ZEB1 and SNAIL [8]. Collectively, mtp53 and MDMX form a cooperative oncogenic network that promotes metastatic potential.

Both PARP and PARG are the enzymes used to fine tune regulation of the DNA damage induced Poly-ADP-ribosylation (PARylation) of proteins [25]. While PARP inhibitors (PARPi) are approved for use in breast cancer patients with homologous recombination deficiencies, like with BRCA1/2 mutations [26], no such approval exists for TNBC with mutations in the *TP53* gene. Given the limited therapeutic options for TNBC, p53 mutation status could serve as an actionable biomarker. We found that combining the PARPi talazoparib (TAL) with the alkylating agent temozolomide (TMZ) induces synergistic cytotoxicity specifically in mtp53, but not wtp53 expressing breast cancer organoids, independent of BRCA status [27]. This synergy associated with the accumulation of replication stress markers that are markers for double-strand DNA breaks and cell death [27]. PARP inhibitors have been shown to increase replication stress by inducing replication gap formation [28]. Inhibiting PARG offers an additional strategy to exploit cancer cell replication stress vulnerabilities. PARG inhibition (PARGi) prevents PAR degradation, causing sustained PAR polymer accumulation, fork collapse, and ssDNA gap formation [29]. Persistent PAR signaling and ssDNA gaps represent distinct liabilities in replication stress-adapted tumors, providing a rationale for combining PARPi or PARGi with DNA-damaging agents to improve treatment outcomes in mtp53-expressing cancers. Despite these insights, how replication stress activates PARP signaling in mtp53 TNBC remains incompletely understood, and whether mtp53 directly establishes a PARP-dependent replication program has not been resolved.

Here, we identify a mechanism in which mtp53, through its CTD, regulates PARP and PAR to sustain DNA synthesis under stress. This mtp53-PARP axis remodels replication stress responses, establishes an unidentified survival pathway, and creates a synthetic vulnerability that can be exploited therapeutically. Remarkably, combining PARP inhibition (TAL) with DNA-damaging therapy (TMZ) in the xenograft mouse models highlights that mtp53 acts as a targetable biomarker for replication stress using modulation of PARylation as an inhibitor of metastasis.

## Results

### Differential DNA Damage Response Pathways in Wild-Type and Mutant p53 Breast Cancer Cells in Response to PARPi (Talazoparib) Plus DNA Damaging Agent (Temozolomide)

DNA damage activates the ATM/ATR-CHK signaling cascade to stabilize wild type p53 (wtp53), and wtp53 activation results in p21-mediated cell cycle arrest to undergo DNA repair or apoptosis [12, 30]. Following repair, wtp53-induced MDM2 provides negative feedback to degrade wtp53 and restore homeostasis, thereby maintaining genomic integrity and cell proliferation [31]. In contrast, mutant p53 (mtp53) disrupts this checkpoint control, converting the DNA damage response (DDR) into a pro-survival, replication stress tolerant state [32]. Previously, we demonstrated that treatment with the combination of talazoparib (TAL) and temozolomide (TMZ) resulted in synergistic cytotoxicity selectively in mtp53-expressing, but not in wild-type p53 breast cancer cells and organoids [2, 27]. Most TNBC have mutations in the *TP53* gene, while estrogen receptor positive (ER+) breast cancers do not. To further examine DDR signaling in this context, we compared wtp53 ER+ MCF7 cells and mtp53 R273H TNBC MDA-MB-468 cells grown on culture dishes, following TMZ, TAL, or combination treatment. The treatment of MCF7 cells with TAL, TMZ, or TMZ + TAL significantly increased p53 protein levels detected in whole cell and chromatin extracts (Fig. 1A, compare lane 1 to 2, 3 and 4 and 9 to 10, 11, and 12). Furthermore, p53 downstream targets p21 and MDM2 were observed to be upregulated when whole cell extracts were compared for vehicle (DMSO) to treatments with the combination treatment inducing the strongest response (Fig. 1A compare p21 and MDM2 in lane 1 to lanes 2, 3, and 4). The inhibition of PARylation by TAL treatment was evident by reduced PARylation in by the context of wtp53 in MCF7 cells and mtp53 in MDA-MB-468 breast cancer cells (see upper PAR panel in Figs 1A and 1B). By contrast, the treatments of MDA-MB-468 cells harboring mtp53 R273H revealed stable mtp53 protein expression in the whole cell extract and chromatin fraction under all conditions (Fig. 1B, see p53 panel) No induction of MDM2 was observed and we detected a reduction in p21 following TMZ + TAL treatment (Fig. 1B, see p21, and MDM2 panels). Furthermore, a pronounced reduction of the oncoprotein MDMX was observed in combination-treated mtp53 MDA-MB-468 cells compared with single-agent treatments (Fig. 1B see MDMX panel). Notably, chromatin bound cleaved PARP (89 kDa cleaved PARP vs 116 kDa Full-length PARP1) and γH2AX, markers of DNA damage and apoptosis were markedly elevated in the combination-treated mtp53 R273H MDA-MB-468 cells (Fig. 1B compare lane 9 to lane 12, but not in wtp53 MCF7 cells (Fig. 1A compare lane 9 to lane 12). Collectively, these results support previous findings that while wtp53 activates a canonical p53-p21-MDM2 repair axis to maintain genomic stability [12, 30], mtp53 cells fail to activate this checkpoint and instead undergo replication stress-associated cytotoxicity, MDMX downregulation, and death in response to combined PARP inhibition and DNA damage.

**Figure 1.**
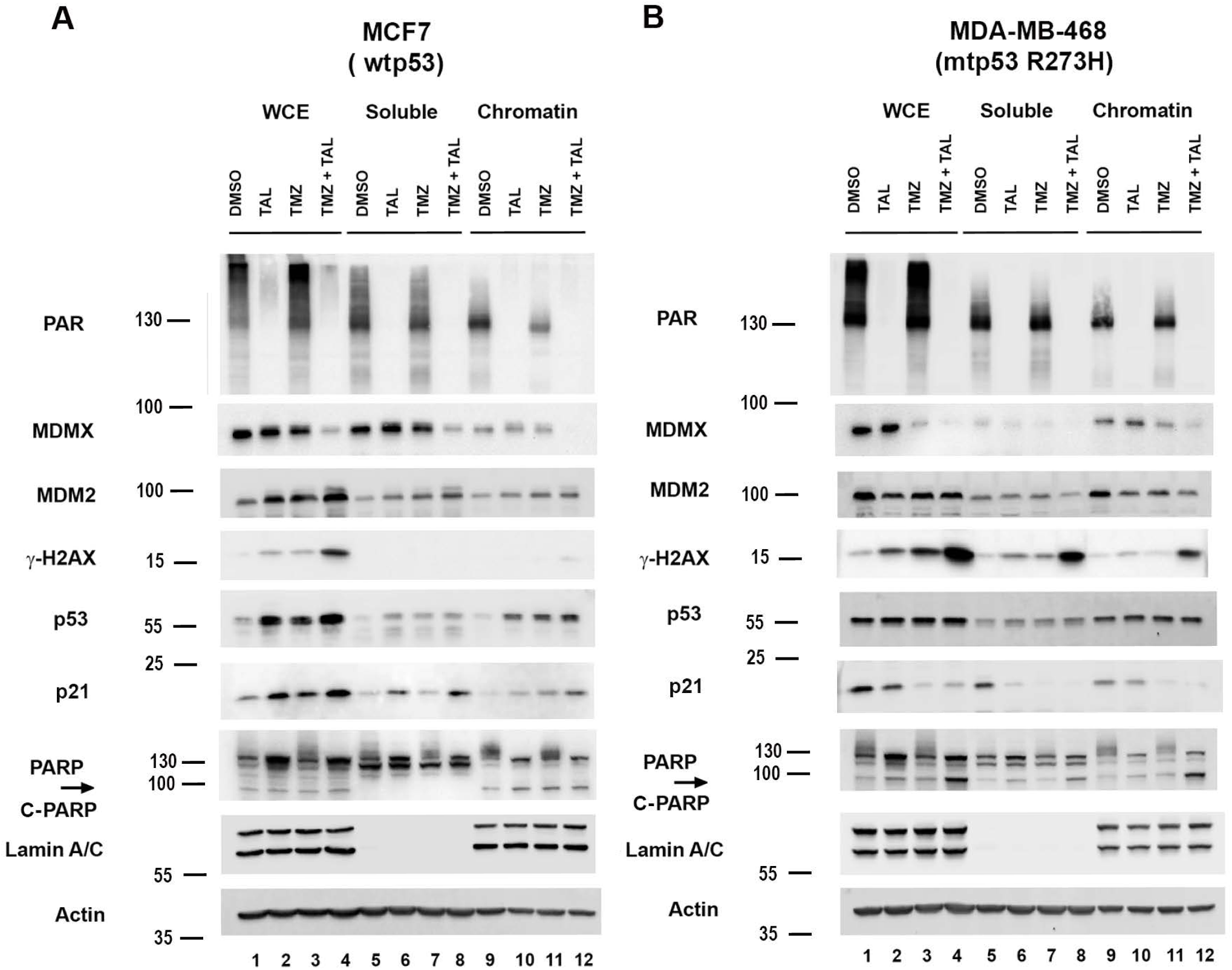
Differential DNA Damage Response Pathways in Wild-Type and Mutant p53 Breast Cancer Cells in Response to PARPi (Talazoparib) Plus DNA Damaging Agent (Temozolomide) Whole cell lysate (WCE), soluble and chromatin fractions were prepared from MCF7 (A) and MDA-MB-468 (B) treated with either vehicle (DMSO), 10 μM talazoparib (TAL), 1 mM temozolomide (TMZ), or combination (TAL + TMZ) for 15 h. Protein levels of PAR,MDMX, MDM2, γ-H2AX, p53, p21 and PARP were determined by Western blot analysis. Representative of two independent experiments.

### Combination Treatment with TMZ plus TAL, Reduces CTCs, Downregulates MDMX and Suppresses Metastatic Signaling in mtp53 Xenografts

While our organoid and cell culture results suggest the TMZ plus TAL combination are effective for mtp53 expressing breast cancer [2, 27], we had yet to test this in a mouse xenograft model. Given the poor prognosis and high metastatic potential of TNBC, largely driven by *TP53* mutations, we evaluated the therapeutic efficacy of combined TMZ plus TAL treatment *in vivo* using MDA-MB-468 xenografts expressing mtp53 R273H. Previous studies in small-cell lung cancer (SCLC) PDX models demonstrated that combination treatment of TMZ and TAL in SCLC patient-derived xenograft (PDX) model significantly suppressed tumor growth and prolonged survival compared to either agent alone [33]. To assess the impact of TMZ + TAL in mtp53 TNBC mouse, we modeled the SCLC PDX protocol [33]. MDA-MB-468 cells were implanted subcutaneously into the flanks of female NOD SCID Gamma (NSG) mice and once tumors reached ∼150 mm³, mice were treated with talazoparib (0.2 mg/kg daily) plus temozolomide (6 mg/kg every 4 days) or vehicle for three cycles following the regimen established by Dr. Rudin’s group (MSKCC). At study endpoint (46 days post implantation) see Supp. Fig. 1A for daily measurement of tumor volume and Supp. Fig. 1B for picture of end point tumor size) circulating tumor cells (CTCs) were quantified as a measure of metastatic dissemination using PE-conjugated anti-human epidermal growth factor receptor (Hu-EGFR) staining and flow cytometry (Fig. 2A). In vehicle controls, MDA-MB-468 xenografts produced an average of 349 CTCs/mL of blood, whereas TMZ plus TAL treatment reduced this number to an average of 25 CTCs/mL, representing a 92.8% reduction. This marked decrease indicated that TMZ plus TAL treatment effectively limited dissemination of mtp53 tumor cells. Consistent with reduced metastatic potential, western blot analysis of whole-cell extracts (WCE) from tumor tissues revealed a significant reduction of MDMX protein levels in TMZ plus TAL treated xenografts compared with controls (Fig. 2B, compare lanes 4, 5 and 6 to lanes 1, 2 and 3). This decrease aligns with our previous observation that MDMX reduction correlates with reduced CTC formation [23]. Downregulation of the EMT transcription factor SNAIL was also observed (Fig. 2B, compare lanes 1, 2 and 3 to lanes 4, 5 and 6). Furthermore, in combination treated tumors, western blot analysis of chromatin fractions demonstrated increased levels of cleaved PARP and γH2AX (Fig. 2C compare lanes 7, 8 and 9 to lanes 10, 11 and 12), indicating replication stress-induced apoptosis. Together, these data demonstrated that combined PARP inhibition and DNA damage effectively target mtp53-driven TNBC by reducing CTC burden and promoting replication stress associated cell death *in vivo*.

**Figure 2.**
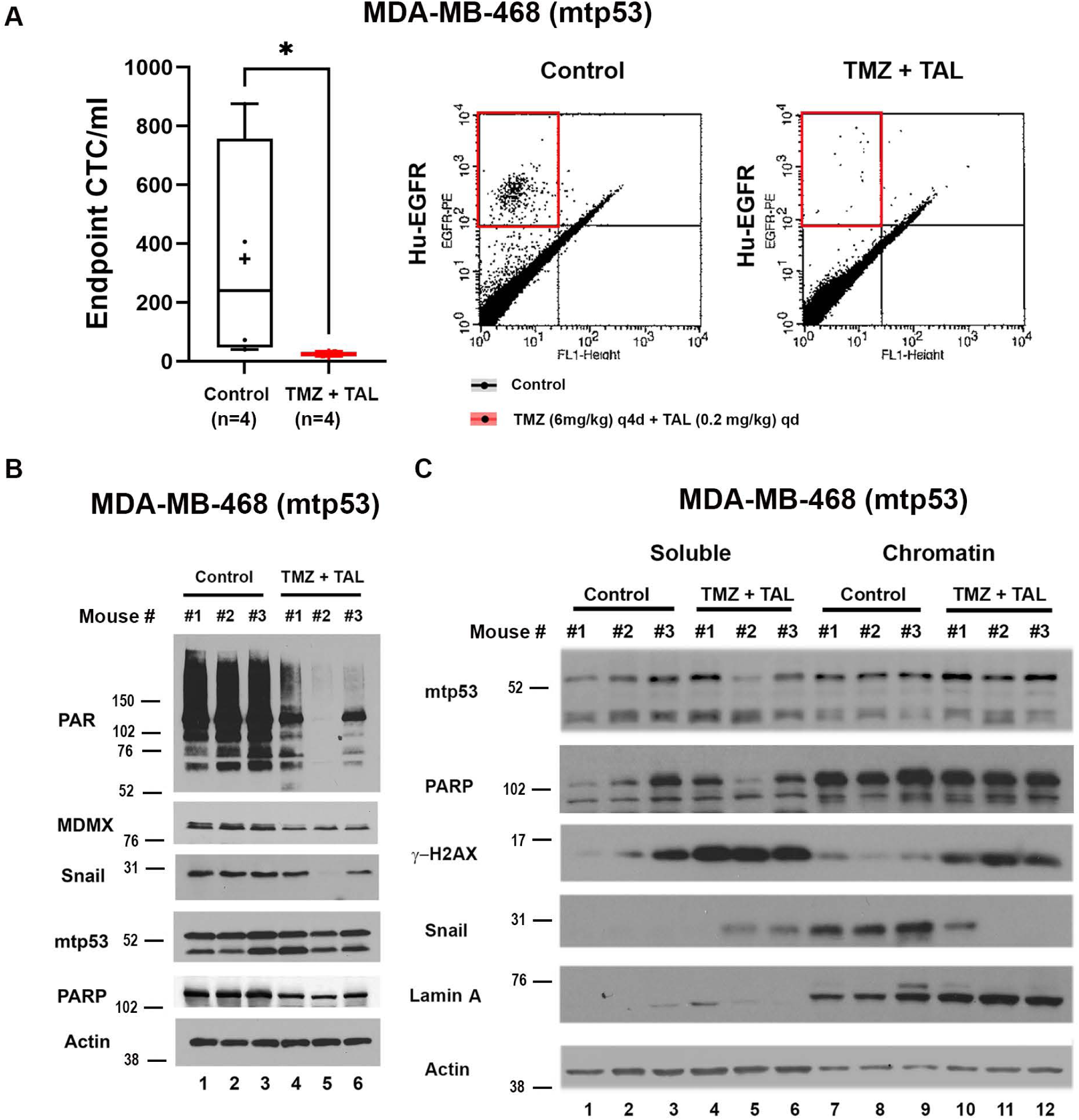
Combination Treatment with TMZ plus TAL, Reduces CTCs, Downregulates MDMX and Suppresses Metastatic Signaling in mtp53 Xenografts. (A) MDA-MB-468 cells were subcutaneous implanted in the flank of female NSG mice. Once tumor volumes reached approximately 150 mm^3^, mice were randomized to treatment via oral gavage either with vehicles (Control), or in combination of talazoparib 0.2mg/kg daily and temozolomide 6 mg/kg every 4 days (TMZ + TAL) treatment group. The CTCs from endpoint MDA-MB-468 xenografts mice were stained with human specific PE-conjugated EGFR antibody and the number of CTCs was determined by flow cytometry. The number of CTCs per milliliter was obtained by dividing the number of positive events by blood volume from individual animals (left). In the box-and-whisker plots, each dot represents one mouse, median values are represented by horizontal lines. + indicated the mean. *P < 0.05, **P < 0.01, ***P <0.001 Mann-Whitney test. Representative fluorescence-activated cell sorting plots showing PE-positive events in Control and TMZ +TAL mouse groups (right). (B) Whole-cell proteins were extracted from NSG mice xenograft tumors and 10 μg of protein was loaded on 10% SDS-PAGE gel. Protein levels of PAR, MDMX, Snail, mtp53 and PARP in endpoint MDA-MB-468 (Control) and MDA-MB-468 (TMZ + TAL) snap-frozen tumors were determined by Western blot analysis. (C) Soluble and Chromatin fractions were prepared from endpoint MDA-MB-468 (Control) and MDA-MB-468 (TMZ +TAL) snap-frozen tumors. Protein levels of p53, PARP, γ-H2AX, and Snail were determined by Western blot analysis.

### Orthotopic Xenograft Model Reveals TMZ Plus TAL Suppresses Metastasis and NF-**κ**B-VEGF Signaling in mtp53 TNBC

To extend our findings from the subcutaneous model showing reduced circulating tumor cells (CTCs) after TMZ + TAL treatment, we next employed an orthotopic implantation strategy to better recapitulate the breast tumor microenvironment. Orthotopic models preserve tumor-stroma interactions and allow spontaneous metastasis, thereby providing a more clinically relevant platform for evaluating therapeutic efficacy [34]. In the wtp53, ER^+^ MCF7 model, cells were implanted into the mammary fat pad of female NSG mice supplemented with estrogen water. Once tumors reached ∼150 mm³, mice were treated with TAL (0.2 mg/kg daily) plus TMZ (6 mg/kg every four days) or vehicle for three cycles. Quantification of lung metastases by immunohistochemistry (IHC) staining of human-specific mitochondrial antigen (Hu-MITO) and QuPath image analysis showed no significant difference between TMZ plus TAL and vehicle-treated groups (Fig. 3A). Consistent with the lack of efficacy, cleaved PARP and γH2AX levels were not increased in the treated animals (Fig. 3B). In contrast, in the mtp53 R273H TNBC MDA-MB-468 orthotopic xenograft model, TMZ + TAL markedly reduced lung metastasis compared with vehicle controls (Fig. 3C, see Supp. Fig. 2B for daily measurement of tumor volume). Hu-MITO IHC and QuPath analysis confirmed significantly decreased metastatic burden. 0.188% Hu-MITO positive cell detection found in vehicle control xenograft lung section, and 0.049% Hu-MITO of cells in TMZ plus TAL xenograft lung section, which is 73.88% reduction. Flow cytometric analysis at the endpoint (58 days post implantation) of CTCs using PE-conjugated anti-human EGFR demonstrated a reduction from an average of 286 CTCs/mL in controls to an average of 57 CTCs/mL in treated mice, representing an ∼80% decrease (Fig. 3D).

**Figure 3.**
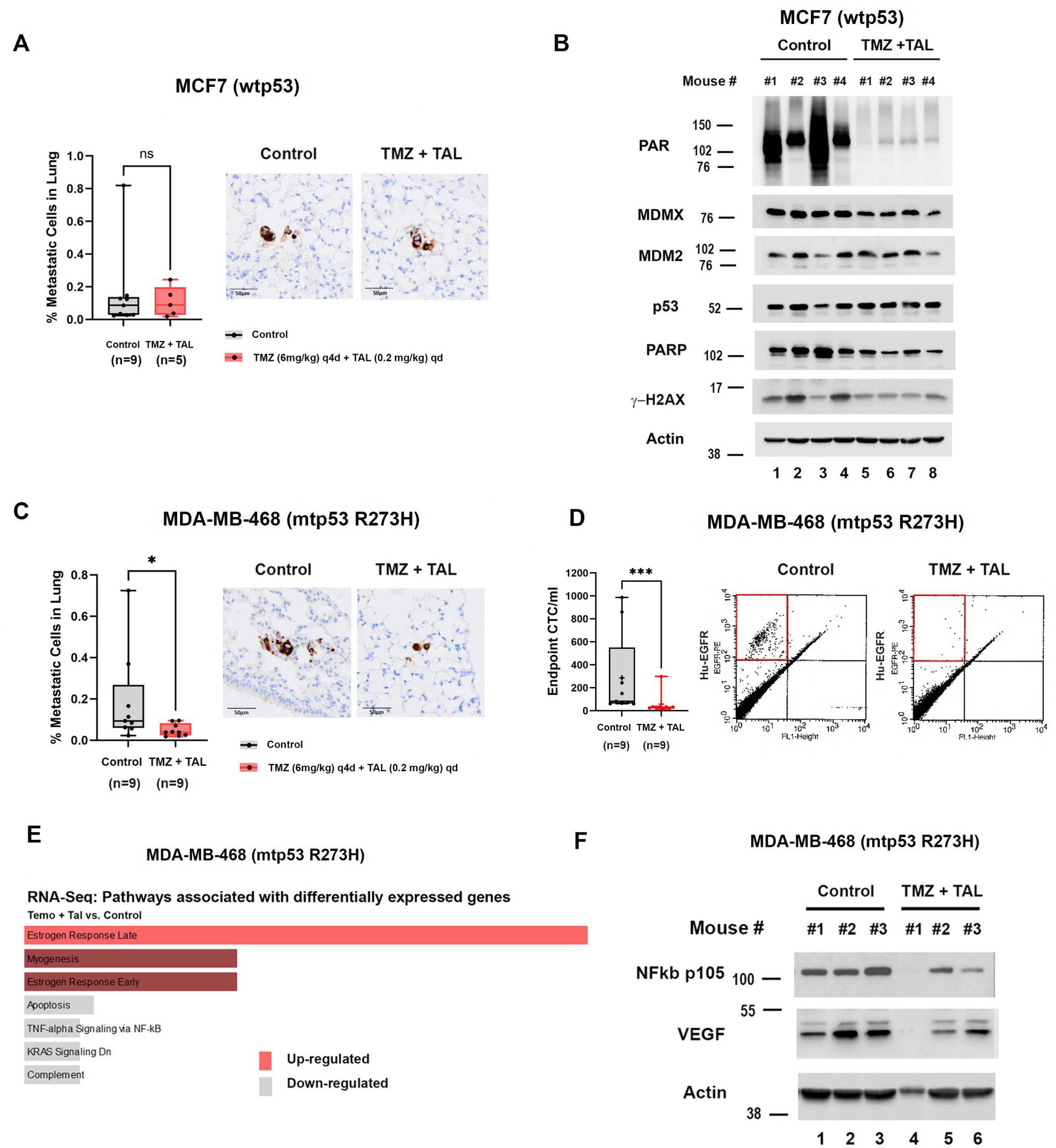
Orthotopic Xenograft Model Reveals TMZ Plus TAL Suppresses Metastasis and NF-κB-VEGF Signaling in mtp53 TNBC. (A) MCF7 cells were orthotopic implanted into the mammary fat pad of female NSG mice. Once tumor volumes reached approximately 150 mm^3^, mice were randomized to treatment via oral gavage either with vehicles (Control), or in combination of talazoparib 0.2mg/kg daily and temozolomide 6 mg/kg every 4 days (TMZ + TAL) treatment group. Lung metastases were measured by IHC staining with human specific mitochondrial antigen on the sections of lungs from orthotopically engrafted MCF7. Quantification of IHC positive cells was assessed with QuPath software (right). (B) Whole-cell proteins were extracted from NSG mice xenograft tumors and 10 μg of protein was loaded on 10% SDS-PAGE gel. Protein levels of PAR, MDMX, MDM2, p53, PARP and, γ-H2AX in endpoint MCF7 Control and TMZ + TAL snap-frozen tumors were determined by Western blot analysis. (C) MDA-MB-468 cells were orthotopic implanted in female NSG mice. Once tumor volumes reached approximately 150 mm^3^, mice were randomized to treatment via oral gavage vehicles (Control), or in combination of talazoparib 0.2mg/kg daily and temozolomide 6 mg/kg every 4 days (TMZ + TAL) treatment group. Lung metastases were measured by IHC staining with human specific mitochondrial antigen on the sections of lungs from orthotopically engrafted MDA-MB-468 Control and TMZ + TAL mouse group. Quantification of IHC positive cells was assesed with QuPath software (right). (D)The CTCs from endpoint MDA-MB-468 xenografts mice were stained with human specific PE-conjugated EGFR antibody and the number of CTCs was determined by flow cytometry. The number of CTCs per milliliter was obtained by dividing the number of positive events by blood volume from individual animals (left). Representative fluorescence-activated cell sorting plots showing PE-positive events in Control and TMZ +TAL mouse groups (right). (E) RNA-seq-based pathway enrichment analysis of MDA-MB-468 (mtp53 R273H) cells treated with temozolomide and talazoparib (TMZ + TAL) compared with control (Method). Gene Set Enrichment Analysis (GSEA) using MSigDB Hallmark gene sets identifies pathways significantly enriched among differentially expressed genes. Up-regulated pathways are shown in red, whereas down-regulated pathways are shown in gray. Enrichment direction reflects normalized enrichment scores derived from genes ranked by differential expression. (F) Whole-cell proteins were extracted from NSG mice xenograft tumors and 10 μg of protein was loaded on 10% SDS-PAGE gel. Protein levels of NF-κB p105 and VEGF in endpoint MDA-MB-468 (Control) and MDA-MB-468 (TMZ + TAL) snap-frozen tumors were determined by Western blot analysis. In the box-and-whisker plots, each dot represents one mouse, median values are represented by horizontal lines. + indicated the mean. *P < 0.05, **P < 0.01, ***P <0.001. Mann-Whitney test.

To identify transcriptional programs modulated by combination therapy, we performed RNA-seq on mtp53 orthotopic xenograft tumors (Fig. 3E). Differential expression and pathway enrichment analyses revealed upregulation of Estrogen Response Early and Late pathways, although typically ERα-independent in TNBC, may involve alternative effectors such as ERβ (ESR2), GPER1, or AR [35, 36]. Importantly, TNF-α/NF-κB signaling, a pathway frequently hyperactivated in TNBC and associated with VEGF-driven angiogenesis and metastasis was significantly downregulated by TMZ + TAL. Transcriptomic profiling showed decreased MDMX and VEGFA mRNA levels (See Supp. Data of RNA-seq data set). Western blot analysis of tumor lysates confirmed that TMZ + TAL reduced full-length NF-κB p105 and concomitantly decreased VEGF protein expression (Fig. 3F), supporting inhibition of NF-κB-dependent VEGF signaling in mtp53 TNBC. Collectively, these results demonstrate that TMZ + TAL treatment suppresses metastasis and angiogenic signaling in mtp53-driven TNBC by downregulating the NF-κB-VEGF axis, thereby impairing tumor vascularization and dissemination.

### TMZ Plus TAL Treatment Inhibits Metastasis of TNBC PDX Models Harboring mtp53 R273H or Lacking p53 Expression

Given the clinical limitations of cell line derived xenografts in recapitulating intratumoral heterogeneity and human-specific signaling interactions, patient-derived xenograft (PDX) models provide a more translationally relevant platform for therapeutic evaluation [37]. PDX tumors maintain the histologic architecture, molecular diversity, and genomic fidelity of the original tumors, enabling robust assessment of treatment response and resistance mechanisms [37]. To validate our findings in a physiologically relevant context, we utilized the PDX models WHIM25, a triple-negative breast cancer (TNBC) model harboring mutant p53 R273H, and WHIM6, a basal-like, ER-negative model that we found does not express p53 [3, 38]. Prior studies demonstrated that WHIM25 exhibits markedly higher PARP and mtp53 protein expression, along with increased PARylated protein levels, compared to WHIM6, reflecting mtp53 enhanced PARP pathway [3]. We evaluated the effect of TMZ plus TAL of the NSG mice xenograft flank implanted metastasis in these PDX models (Fig. 4A & 4B). (see Supp. Fig. 3A for daily measurement of tumor volume and Supp. Fig. 3B for picture of end point tumor size). At study endpoint (47 days post implantation of WHIM25 and 36 days post implantation of WHIM6), CTCs were quantified using PE-conjugated anti-human EGFR staining and flow cytometry. In vehicle-treated controls, the WHIM25 PDX generated an average of ∼60 CTCs/mL of blood, whereas TMZ plus TAL treatment reduced CTCs to ∼30 CTCs/mL, representing a 50% reduction (Fig. 4A). WHIM6 PDX produced ∼45 CTCs/mL in controls, reduced to ∼18 CTC/mL with combination treatment, representing a 60% reduction (Fig. 4B). These data indicate that TMZ plus TAL effectively limited either blood dissemination or blood stream cancer cell survival.

**Figure 4.**
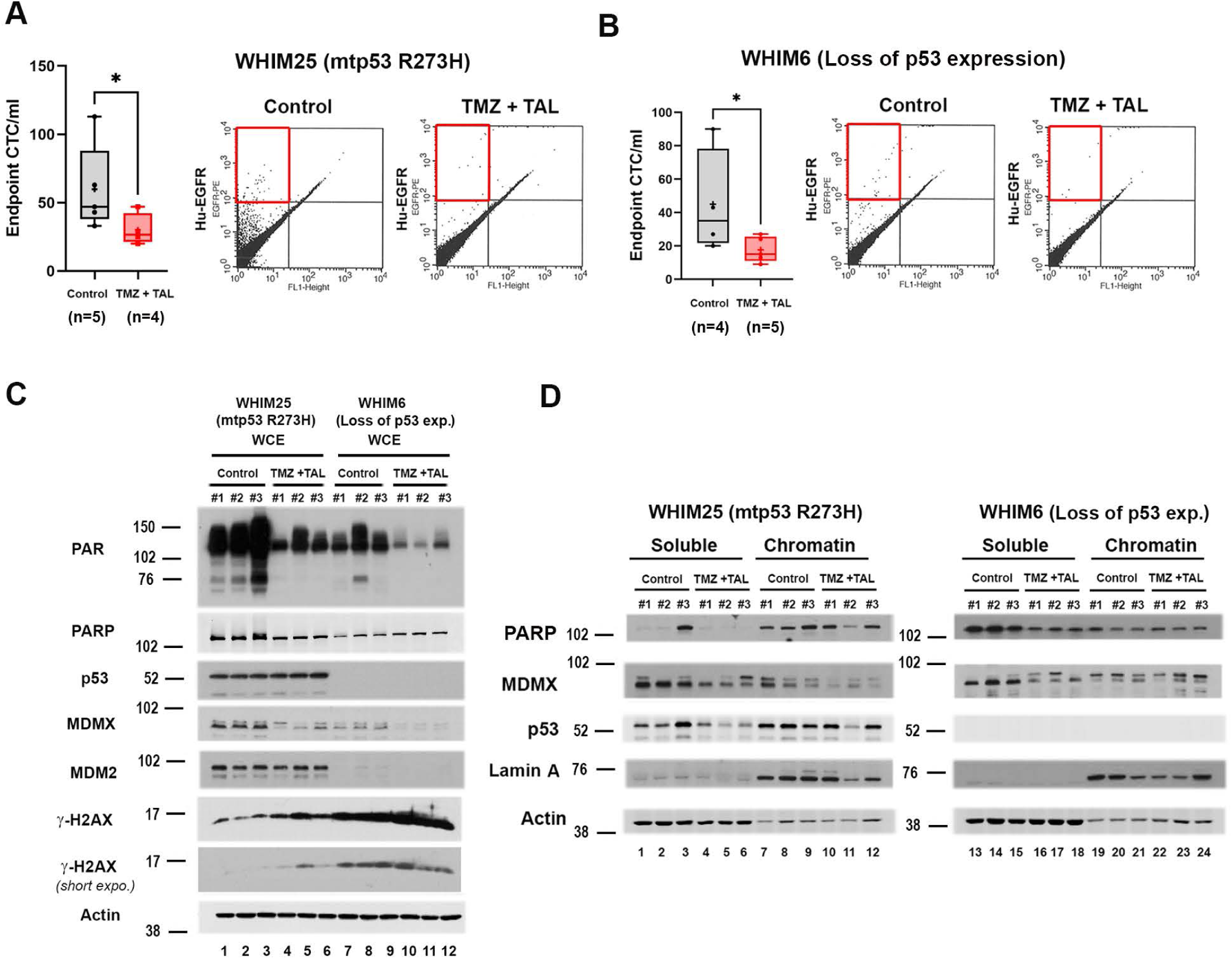
TMZ plus TAL Treatment Inhibits Metastasis of TNBC PDX Models Harboring mtp53 R273H or Lacking p53 Expression. (A) Breast cancer PDX model WHIM25 (A) WHIM6 (B) cells were subcutaneous implanted in the flank of female NSG mice. Once tumor volumes reached approximately 150 mm^3^, mice were randomized to treatment via oral gavage either with vehicle (Control), or in combination of talazoparib 0.2mg/kg daily and temozolomide 6 mg/kg every 4 days (TMZ +TAL) treatment group. The CTCs from WHIM25 or WHIM6 Control or TMZ + TAL mice were stained with human specific PE-conjugated EGFR antibody, and the number of CTCs was determined by flow cytometry. The number of CTCs per milliliter was obtained by dividing the number of positive events by blood volume from individual animals. (C) Whole cell protein levels of PAR, PARP, p53, MDMX, MDM2 and γ-H2AX in endpoint Control and TMZ + TAL snap-frozen tumors were determined by Western blot analysis. Whole-cell proteins were extracted from NSG mice xenograft tumors and 10 μg of protein was loaded on 10% SDS-PAGE gel. (D) Soluble and Chromatin fractions were prepared from endpoint WHIM25 or WHIM6 Control and TMZ +TAL snap-frozen tumors. Protein levels of PARP, MDMX and p53 were determined by Western blot analysis. In the box-and-whisker plots, each dot represents one mouse, + indicated the mean. *P < 0.05, **P < 0.01, ***P <0.001, Mann-Whitney test.

We addressed biomarkers in the primary tumors to evaluate markers of cell death and metastasis. Consistent with reduced metastatic potential, immunoblot analysis of whole-cell extracts (WCE) revealed a significant reduction of MDMX protein levels in TMZ plus TAL treated tumors from both WHIM25 and WHIM6 (Fig. 4C). Notably, despite wtp53 status in WHIM6, p53 was not detectable and no transcriptional activation was observed of the canonical target gene MDM2 in response to DNA damage. Analysis of chromatin fractions demonstrated increased DNA damage in combination-treated WHIM25 tumors, indicated by increased γH2AX (Fig. 4D). Reduction of MDMX was observed in both soluble and chromatin fractions in WHIM25, whereas in WHIM6, MDMX reduction was primarily in the soluble fraction. Collectively, these findings suggest that TMZ plus TAL treatment suppresses metastatic potential and induces replication stress-associated cell death preferentially in mtp53 TNBC, while also limiting dissemination in tumors lacking functional p53.

### CRISPR-Mediated Deletion of the Mutant p53 C-terminus Reduces Tumor Growth and Metastatic Dissemination in a TNBC Orthotopic Xenograft Model

The CRISPR/Cas9-mediated genome edited MDA-MB-468 cells with the C-terminal deletion mutant p53 R273H(Δ347-393) and a frameshift mutant p53 R273Hfs387 markedly reduces cell proliferation [4]. The deletion of the C-terminus of mutant p53 R273H also reduces sensitivity to TMZ plus TAL treatment [5]. In addition, the C-terminal region of mtp53 plays a critical regulatory role in mediating an interaction with PARP and poly(ADP-ribose) [5]. To determine whether this interaction contributes to tumor progression and metastatic potential, we tested these cells in an orthotopic xenograft model to assess the functional significance of the mtp53 C-terminus for tumor formation and metastasis.

We validated that they grew well in 2D tissue culture and had the expected p53 protein size after CRISPR deletion. They also showed variation in mtp53, PARP, PAR, and MDMX expression levels (Fig. 5A). The C-terminally truncated mtp53 R273H(Δ347-393) was detected exclusively in the soluble fraction, with no detectable chromatin association (Fig. 5A compare lane 5 to lane 8), while mtp53 protein levels were markedly reduced in R273Hfs387 cells (Fig. 5A compare lane 3 to lane 1). Notably, chromatin-bound PARP also decreased in R273Hfs387 cells, consistent with our previous findings implicating mtp53-mediated PARP chromatin binding (Fig. 5A compare lane 9 to lane 7). Interestingly, MDMX protein levels were reduced in R273Hfs387 cells across WCE, soluble, and chromatin fractions, and in the chromatin fraction of R273HΔ347-393 cells, compared to mtp53 R273H MDA-MB-468 controls. Moreover, R273Hfs387 cells exhibited a distinct PARylation pattern, characterized by slower-migrating PAR-modified species, suggesting altered PARP signaling dynamics in the absence of full-length mtp53.

**Figure 5.**
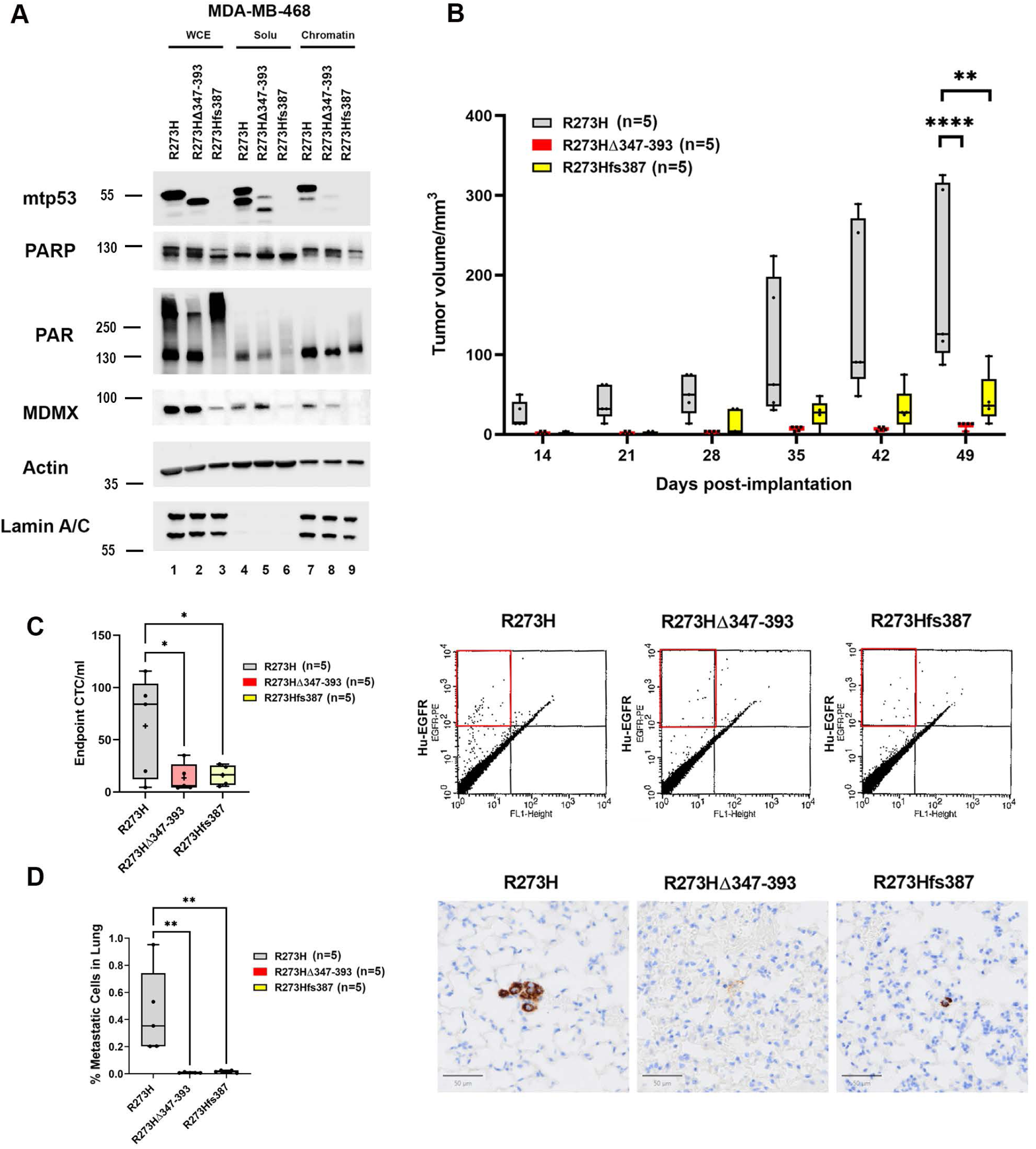
CRISPR-Mediated Deletion of the Mutant p53 C-terminus Reduces Tumor Growth and Metastatic Dissemination in TNBC Orthotopic Xenograft Model. (A) Whole cell lysate (WCE), soluble and chromatin fractions were prepared from MDA-MB-468 cells which express mtp53 R273H, and clones (R273HΔ347-393 or R273Hfs387) generated via CRISPR/Cas9 gene editing. Protein levels of mtp53 PARP, PAR and MDMX were determined by Western blot analysis. (B) MDA-MB-468 cells which express mtp53 R273H, and CRISPR clones R273HΔ347-393 or R273Hfs387 were orthotopically implanted into the mammary fat pad of female NSG mice. Primary tumor volumes from each mouse were measured using calipers. In the box-and-whisker plots, each dot represents one mouse, + indicated the mean. Endpoint tumor volumes (transformed in Log) were performed with ordinary one-way ANOVA test, *P < 0.05, **P < 0.01, ***P <0.001. (C) The CTCs from engrafted NSG mice were stained with human specific PE-conjugated EGFR antibody and the number of CTCs was determined by flow cytometry. The number of CTCs per milliliter was obtained by dividing the number of positive events by blood volume from individual animals (left) In the box-and-whisker plots, each dot represents one mouse, + indicated the mean. *P < 0.05, **P < 0.01, ***P <0.001, ordinary one-way ANOVA. Representative fluorescence-activated cell sorting plots showing PE-positive events in different mouse groups (right). (D) Lung metastases were measured by IHC staining with human specific mitochondrial antigen on the sections of lungs from orthotopically engrafted MDA-MB-468 mtp53R273H, R273HΔ347-393 or R273Hfs387 cells. Quantification of IHC positive cells was accessed with QuPath software (left) In the box-and-whisker plots, each dot represents one mouse, + indicated the mean. *P < 0.05, **P < 0.01, ***P <0.001, ordinary one-way ANOVA. Representative of human specific mitochondrial antigen IHC showing positive staining in different mouse groups (right).

Next, we investigated if the C-terminus of mutant p53 contributed to tumor development *in vivo*. A total of 5 × 10 MDA-MB-468 cells expressing mtp53 R273H, mtp53 R273H(Δ347-393), or mtp53 R273Hfs387 were implanted into the mammary fat pad of female NSG mice. While both mtp53 R273H(Δ347-393) and mtp53 R273Hfs387 can grow in culture, the xenografts exhibited strikingly significantly reduced tumor growth compared to parental MDA-MB-468 tumors. At the study endpoint, we observed an average 93.98% reduction in tumor volume in mtp53 R273H(Δ347-393) xenografts and an average 77.12% reduction in mtp53 R273Hfs387 xenografts (Fig. 5B 49 days post-implantation, see Supp. Fig. 4 for picture of end point tumor size). We evaluated the impact of the mtp53 C-terminus on metastatic dissemination (Fig. 5C-D). At the study endpoint, circulating tumor cells (CTCs) were quantified by PE-conjugated anti-human EGFR staining and flow cytometry. In parental MDA-MB-468 xenografts, an average of ∼63.2 CTCs/mL of blood was detected, whereas mtp53 R273H(Δ347-393) xenografts showed a reduction to ∼13.4 CTCs/mL (78.7% reduction), and mtp53 R273Hfs387 xenografts showed ∼16.1 CTCs/mL (74.5% reduction) (Fig. 5C). Furthermore, IHC and QuPath analysis confirmed a significant decrease in lung metastatic burden in both mtp53 C-terminal deletion and frameshift xenografts (Fig. 5D). 0.449% human-specific mitochondrial antigen IHC positive cell detection was found in mtp53 R273H xenograft lung sections, 0.009% of cells in mtp53 R273H(Δ347-393) xenografts, and 0.017% of cells in mtp53 R273Hfs387 xenografts, corresponding to 98.09% and 96.13% reductions, respectively, compared to mtp53 R273H lung sections. In conclusion, *in vivo* CRISPR-mediated deletion of the mtp53 C-terminal region (Δ347-393) or frameshift mutation (R273Hfs387) in TNBC xenografts resulted in markedly reduced tumor growth, circulating tumor cells, and lung metastasis, demonstrating that the C-terminus of mtp53 is critical for its tumor-promoting and metastatic functions when tested in a xenograft model.

### Replication Stress and PAR Signaling Are Sustained in mtp53 but Not in C-Terminal-Deleted mtp53 R273H Cells

To dissect if full-length gain of function mutant p53 R273H modulates replication stress and PAR signaling, we employed PARG inhibition (PARGi) to evaluate PAR dynamics and the replication stress response. This enabled us to test if mtp53 R273H and C-terminal-deleted mtp53 R273H (Δ347-393) contributed to PAR homeostasis and replication fork stability. Inhibition of PARG disrupts PAR turnover, resulting in persistent single-stranded DNA (ssDNA) gaps during DNA replication [29]. To quantitatively compare fork progression and ssDNA gap formation in the different isogenic cell lines, we used the DNA fiber combing assay combined with S1 nuclease treatment, which specifically digests regions of ssDNA to reveal replication stress gaps.

Both mtp53 R273H and C-terminal-deleted mtp53 R273H (Δ347-393) MDA-MB-468 cells were sequentially labeled with CldU (30 min) followed by IdU (60 min) in the presence or absence of PARGi. After labeling, cells were lysed, and genomic DNA was treated with or without S1 nuclease for 30 minutes prior to fiber combing. IdU track lengths were then measured to assess replication fork progression (schematic and representative fibers shown in Fig. 6A; quantification in Fig. 6B). In mtp53 R273H cells, both S1 and PARG inhibition significantly reduced IdU track length. Compared to the control average (22.30 μm), track length decreased to 15.69 μm with S1 treatment (29.6% reduction) and 14.80 μm with PARGi (33.6% reduction). The combination of PARGi + S1 further shortened IdU tracks to 9.81 μm, representing a 56.0% reduction relative to control. These results indicate that mtp53 R273H cells accumulate extensive ssDNA gaps and undergo severe replication fork degradation when PARG activity is blocked, consistent with elevated replication stress and fork instability. In contrast, C-terminal-deleted mtp53 R273H (Δ347-393) cells showed minimal change across all conditions (Control: 16.67 μm; S1: 17.07 μm [+2.4%]; PARGi: 16.85 μm [+1.1%]; PARGi + S1: 16.23 μm [-2.6%]). The absence of IdU shortening suggests that loss of the CTD confers resistance to PARGi and limits ssDNA gap accumulation. Under control conditions, mtp53 R273H cells exhibited a 25.2% longer IdU track length than C-terminal-deleted mtp53 R273H cells (22.30 μm vs 16.67 μm), suggesting that full-length mtp53 accelerates fork progression. Together, these data demonstrate that the C-terminal domain of mtp53 R273H is essential for promoting PARG-dependent replication stress and fork instability.

**Figure 6.**
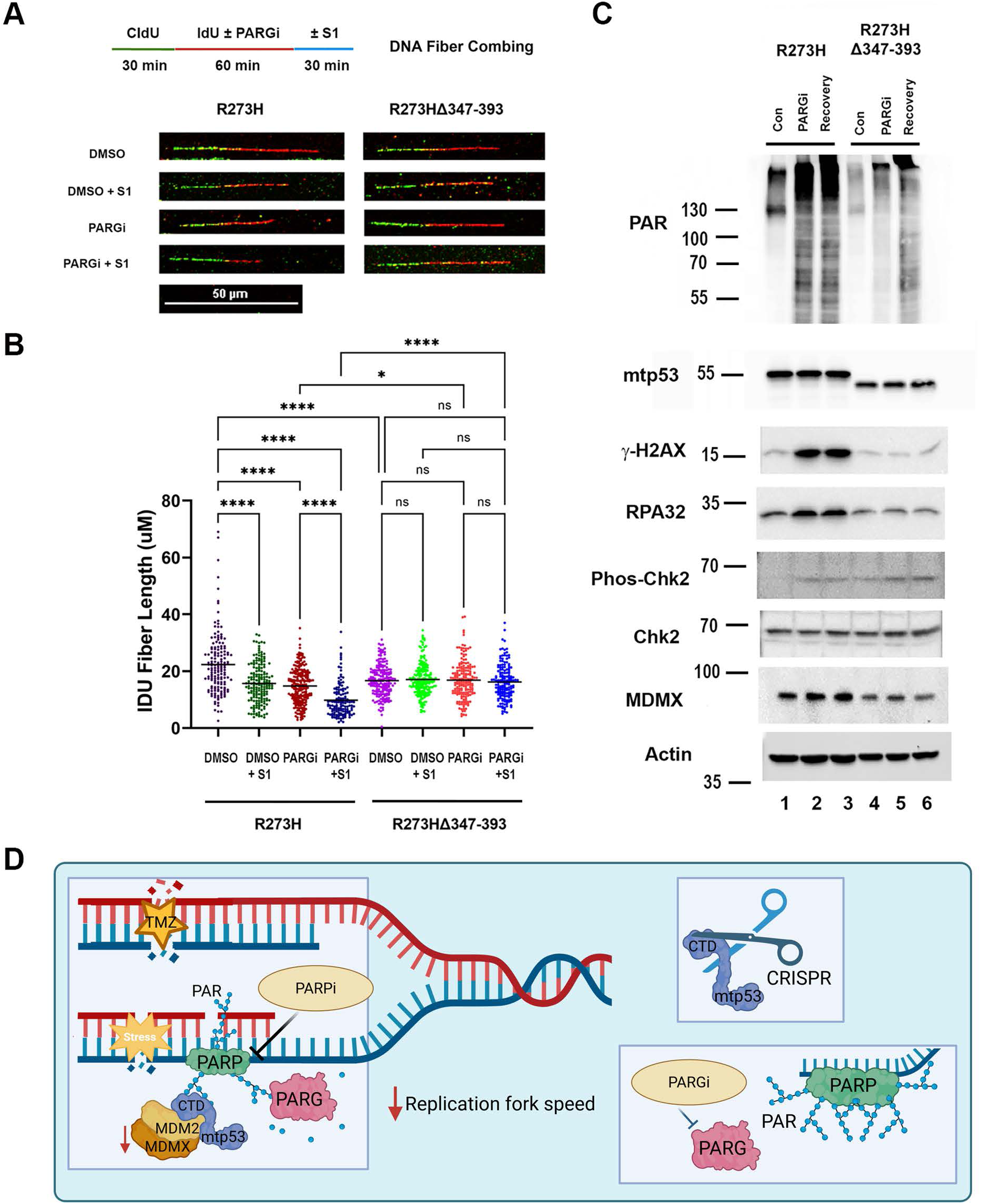
Replication Stress and PAR Signaling Are Sustained in mtp53 but Not in C-Terminal-Deleted mtp53 R273H Cells. (A) Top: Schematic of DNA fiber combing assay with CldU/IdU pulse-labeling protocol and treatment with PARGi, followed by S1 nuclease treatment. Bottom: Representative DNA fiber images of MDA-MB-468 cells with mtp53 R273H, and mtp53 (R273HΔ347-393 with vehicle (DMSO) or PARGi with or without S1 nuclease treatment. (B) Quantification of IdU tracts after 10 μM PARGi treatment with or without S1 nuclease. Each dot represents one fiber. Around 130-220 fibers were analyzed for each condition. Mean values are represented by horizontal lines. ns., not significant *P < 0.05, **P < 0.01, ***P <0.001, ****P < 0.0001, two-way ANOVA test followed by Tukey’s post-hoc test. Representative of two independent experiments. (C) Whole cell protein levels of PAR, mtp53, γ-H2AX, RPA32, Phos-Chk2, Chk2 and MDMX were determined by Western blot analysis. Representative of two independent experiments. (D) Model for mutant p53-PARP axis enables replication stress tolerance and drive cancer metastasis, created in BioRender.

Consistent with these findings, biochemical analysis by western blot further confirmed replication stress activation as indicated by increased γH2AX (Fig. 6C, compare lane 2 to lane 1). Treatment of mtp53 R273H cells with PARGi for 60 minutes followed by 24-hour recovery led to accumulation of PAR, increased γH2AX, phospho-Chk2, and RPA32, indicating replication stress and DNA damage signaling triggered by ssDNA gap persistence. In contrast, C-terminal deleted mtp53 R273H (Δ347-393) cells, which displayed minimal IdU shortening, showed markedly attenuated PAR, γH2AX, p-Chk2, and RPA responses following PARGi. These findings establish the C-terminal region of mtp53 as a critical mediator of replication stress tolerance. Full-length mtp53 R273H promotes sustained replication stress, maintaining proliferation despite persistent ssDNA gaps. In contrast, deletion of the CTD abolishes this adaptive response, slowing replication and reducing tumor formation. This underscores the role of the mtp53 C-terminal domain in maintaining DNA replication fork stability under the stressful conditions encountered in tumors.

## Discussion

This study uncovers the critical role of mutant p53 (mtp53) in coordinating PARP-dependent replication stress tolerance and metastatic progression in TNBC. Our findings revealed that the therapeutic outcome of combined temozolomide (TMZ) and talazoparib (TAL) treatment to reduce tumor burden and metastasis occurred in mtp53 expressing, but not wtp53 expressing breast cancer. Following TMZ plus TAL treatment mutant p53 cells accumulated double-strand breaks indicated by increased γH2AX and showed marked MDMX reduction. Significantly, TMZ plus TAL treatment reduced circulating tumor cells (CTCs) and lung metastases of mutant, but not wtp53 expressing cells. In mtp53 R273H MDA-MB-468 cells, xenograft, and mtp53 R273H WHIM25 PDX models, MDMX protein levels were consistently downregulated after TMZ plus TAL treatment, and this MDMX destabilization correlated with increased replication stress-associated cytotoxicity and reduced metastatic fitness. Our findings further demonstrated that the C-terminal domain (CTD) of mtp53 acts as a structural and regulatory hub linking PARP and PAR, maintaining replication fork progression in the presence of ssGAPs. CRISPR-mediated CTD deletion disrupted mtp53 R273H and PAR chromatin binding, slowed replication dynamics, and markedly inhibited tumor growth and metastasis. This establishes the mtp53 CTD as a key determinant of replication stress adaptation and a potential therapeutic vulnerability (Fig. 6D).

Despite being the most frequently mutated gene in human cancer, mutant p53 remains an “undruggable” target due to its nuclear localization and absence of defined ligand-binding pockets [39]. Consequently, we have shifted our therapeutic focus toward targeting gain of function of mtp53 PARP addiction, replication stress tolerance, and DNA repair defects.

In wild-type p53 (wtp53) breast cancer cells, DNA damage triggers canonical p53-p21-MDM2 signaling, that induces cell cycle arrest and DNA damage repair [40]. By contrast, mtp53 cells fail to activate this checkpoint, continuing DNA synthesis under stress. It has been shown that PARP recruits MRE11 to stabilize stalled forks, maintaining replication recovery capacity [16, 17]. Our study further indicated that mtp53 maintains fork stability, thereby facilitating continuous proliferation under a DNA gap stressed fork accumulation and may do so using the C-terminus to coordinate PARP. Mechanistically, mtp53 stabilizes chromatin-bound PARP, forming an oncogenic replication complex that supports fork progression. In addition, mtp53 interacts with key fork remodeling and restart factors-such as MRE11 [41], suggesting mtp53 could stabilize stalled forks and enable replication under stress. Perhaps mtp53 helps PRIMPOL, SMARCAL1 and ZRANB3 and balance repriming versus fork reversal pathways. Deletion of the CTD abolished mtp53 chromatin binding and PARP recruitment, resulting in defective fork progression because of ineffective fork restart. These findings reveal that full-length mtp53 coordinates a PARP/PAR that allows cells to proliferate despite unresolved DNA lesions, while CTD loss disassembles this replication stress tolerance network.

Our results establish a direct link between mtp53-PAR signaling and metastatic competence. TMZ plus TAL treatment reduced MDMX, NF-κB, and VEGF expression in mtp53 tumors, correlating with decreased CTC numbers and metastatic lung foci. These results suggest that TMZ plus TAL disrupts oncogenic signaling networks sustaining metastasis in mtp53 cells. Importantly, these effects extended to the p53-deficient WHIM6 PDX model, suggesting a common vulnerability exploitable by PARP inhibition combined with DNA damaging agents. Previously, we discovered that MDMX promotes CTC formation in mtp53 MDA-MB-231 xenograft model [23] and mtp53 can promote metastasis via NF-κB-VEGF activation [8, 42, 43].Reduction of MDMX and targeting mtp53 likely explains the profound reduction in CTCs and lung metastases seen in mtp53 xenograft models. Together, our results define two mtp53-driven oncogenic programs that are suppressed by TMZ plus TAL: (i) adaptation to replication stress through PARP engagement, and (ii) promotion of metastatic dissemination via NF-κB-VEGF signaling. Disrupting this mtp53-PARP axis offers a mechanistic basis for targeting TNBC subtypes that depend on replication-stress tolerance. Moreover, these findings support mtp53 as a predictive biomarker for PARP-based combination therapies, expanding PARP inhibitor clinical utility beyond BRCA-mutant cancers.

The reduction of CTCs and lung metastases in TMZ plus TAL treated mice underscores how replication stress-targeted therapy can disrupt the systemic dissemination process. High CTCs often mirror the most therapy-resistant tumor clones, and their survival is reinforced by interactions with immune and stromal cells [44]. It has been shown that neutrophil-assisted CTC clusters promote metastatic seeding by shielding tumor cells from immune clearance through neutrophil extracellular trap (NET) formation [45]. Notably, mtp53-driven TNBC exhibits strong NF-κB and cytokine activity [46]. Therefore, therapeutics that block CTC-neutrophil adhesion or degrade NET scaffolds could synergize with TMZ plus TAL by eliminating CTCs. Mutant p53 induces chromosomal instability (CIN) and a cytosolic DNA response that activates cGAS-STING while simultaneously reprograming cytokine output to evade immune surveillance [47]. Prospectively, combining PARP inhibitor with immunotherapy represents a promising avenue. The DNA damage and cytosolic DNA accumulation induced by TMZ plus TAL may trigger cGAS-STING-interferon signaling, enhancing antigen presentation and T-cell infiltration. Immune checkpoint blockade (anti-PD-1/PD-L1) or STING agonists could amplify these effects, converting “immune-cold” mtp53 tumors into immune-responsive lesions.

Beyond TMZ, several FDA-approved DNA-damaging agents, including carboplatin, cisplatin, cyclophosphamide, doxorubicin, epirubicin, topotecan, and capecitabine can synergize with PARP inhibition by activating replication stress [48]. ATR and ATM inhibitors further enhance this strategy by disabling the replication stress checkpoint, forcing fork collapse and apoptosis in mtp53 cancers [49].

In addition, we observed a preliminary finding that the combination of the dual MDM2/MDMX inhibitor ALRN-6924 (ALRN) with TAL to block PARP was not synergistic (Supplementary Fig. 5). In wtp53 MCF7 cells, ALRN or ALRN plus TAL increased p53, MDM2, and MDMX levels, consistent with p53 activation and autoregulatory feedback, but no PARP cleavage was observed (Supplementary Fig. 5A, lanes 3 and 4). In mtp53 R273H MDA-MB-468 cells, neither MDM2/MDMX levels nor DNA damage markers changed after ALRN plus TAL treatment (Supplementary Fig. 5B). Neither MCF7 cells nor MDA-MB-468 cells demonstrated reduced mitochondrial activity when blocking both PARP activity and complex formation of MDM2 with p53 (Supplementary Fig. 5C) Patient-derived tumor organoids (PDTOs) harboring mtp53 R248W or R213stop mutations showed no synergy, and rather evidence of antagonism when combined. These results suggest that direct MDM2/MDMX blockade may disrupt mutant p53 interaction with MDM2 that aids in the PARPi-induced cytotoxicity (Supplementary Fig. 5D). Therefore, while MDM2/MDMX inhibitors can potentiate activation in wtp53 tumors, they may counteract PARPi efficacy in mtp53 contexts.

Unlike catalytic PARP inhibitors and PARP trapping drugs, PARP PROTACs induce proteasomal degradation of PARP1/2, preventing chromatin rebinding and achieving potent DNA repair suppression [50, 51]. Given mtp53’s reliance on PARP signaling for fork stabilization, selective PARP degradation may disrupt this oncogenic complex. However, if PARP trapping is critical for synthetic lethality, then degraders may be ineffective. The PARG inhibition (PARGi), conversely, blocks PAR hydrolysis, locking PARP1 in an inactive trapped state and exhausting NAD⁺ pools, potentially synergizing with replication stress-inducing agents in mtp53 TNBC. Thus, the PARGi strategy may be the most potent synthetic lethal strategy.

Collectively, our data define the mtp53 C-terminal domain as a regulatory hub coupling replication stress tolerance with metastatic signaling. Targeting the mtp53-PARP axis through combination of TMZ plus TAL provides a mechanistic strategy to overcome mtp53’s undruggable nature. By exploiting replication stress vulnerabilities and simultaneously targeting CTC-mediated metastatic dissemination, the combination of a PARP inhibitor or a PARG inhibitor with a DNA-damaging agent holds strong therapeutic potential for improving outcomes in mutant p53-driven TNBC.

## Materials and methods

### Cell culture and patient derived xenograft (PDX) cell lines

Human breast cancer cell lines MDA-MB-468 and MCF7 were purchased from ATCC. Cells were authenticated and tested for mycoplasma (Genetica DNA Laboratories). Cells were maintained at 5% CO2 in DMEM (Invitrogen) with 50 U/ml penicillin, 50 μg/ml streptomycin (Mediatech) and supplemented with 10% FBS (Gemini) in 37°C humidified incubator. PDX lines were generated at Washington University from breast cancer patients [38].

### Drugs

Talazoparib (Cat# S7048) and temozolomide (Cat# S1237) were purchased from Selleckchem. Poly(ADP-ribose) glycohydrolase (PARG) inhibitor was purchased from MedChemExpress (MCE) (Cat# HY-133531). Drugs were diluted and utilized according to manufacturers’ instructions.

### Antibodies

Proteintech: Cat# 84534-5-RR MDMX/MDM4 Rabbit; Cat# 10442-1-AP P53 Rabbit PolyAb; Cat# 10355-1-AP P21 Rabbit PolyAb; Cat# SA00001-2 HRP-conjugated Goat Anti-Rabbit IgG(H+L), Cell Signaling: Cat# 9718S Phospho-Histone H2A.X (Ser139) (20E3) Rabbit mAb; Cat# 89190S Poly/Mono-ADP Ribose (D9P7Z) Rabbit mAb; Cat# 4717S NF-κB1 p105 Antibody; Cat# 7074S Anti-rabbit IgG, HRP-linked Antibody; Cat# 3879S Snail (C15D3) Rabbit mAb; Cat# 2661S Phospho-Chk2 (Thr68) Antibody; Cat# 2662S Chk2 Antibody, Abcam: Cat# AB6326-1001 Anti-BrdU antibody; Cat# AB259265-1001 Anti-MDM2 antibody; Cat# AB191217-1001 Anti-PARP1 antibody; Cat# AB46154-1001 Anti-VEGFA antibody.

### Whole cell lysis and chromatin fractionation

Whole cell lysis was carried out using RIPA buffer (0.1% SDS, 1% IGEPAL NP-40, 0.5% Deoxycholate, 150 mM NaCl, 1 mM EDTA, 0.5 mM EGTA, 50 mM Tris-Cl pH 8.0, 1mM PMSF, 8.5 μg/ml Aprotinin, 2 μg/ml Leupeptin and phosphatase inhibitor cocktail) on ice for 30 min and with occasional vortexing. Samples were sonicated three times for 30 seconds of pulse, followed by 30 seconds of rest on ice/water at 98% amplitude. Samples were centrifuged at 15,700 g at 4°C for 20 min and supernatant was saved. Cell fractionations were prepared using Chromatin Extraction Kit from Abcam (Cat# ab117152) according to the manufacturer’s protocol. Cells were first lysed in Lysis Buffer (1X) containing proteinase and phosphatase inhibitors on ice for 10 min, vortexed vigorously for 10 sec and centrifuged at 2,300 g for 5 min. The supernatant was saved as the soluble fraction and extraction buffer was added to the pellet to resuspend it by pipetting up and down and incubating the sample on ice for 10 min, with occasional vortex mixing. The sample was resuspended and sonicated two times for 30 seconds pulse/followed by 30 seconds of rest at 98% amplitude on ice to increase chromatin extraction. Then, the sample was centrifuged at 15,700 g at 4°C for 10 min. The supernatant was transferred to a new vial and Chromatin Buffer was added at a 1:1 ratio and saved as chromatin bound protein. Protein concentration was determined by Protein Bradford by NanoDrop One (Thermo Scientific).

### Western blot

Proteins were run on either 8%, 10% or 15% SDS-PAGE to separate samples followed by electro-transfer onto PVDF membrane (GE). The membranes were blocked with 5% non-fat milk (Biorad) in 1X Tris-Buffered Saline (TBS)/0.1% Tween-20 following incubation with primary antibody overnight at 4 °C. The membranes were washed 3 times of 1X TBS/0.1% Tween-20 and incubated with secondary antibody for 1 h at room temperature. The signal was detected by chemiluminescence with Super Signal Kit (Pierce) and autoradiography with Hyblot CL films (Denville Scientific) or ChemiDoc Imaging System (Bio-Rad).

### Ethics

All animal experiments were done in accordance with protocols approved by the Institutional Animal Care and Use Committees (IACUC) of Hunter College, Weill Cornell Medical College, and Memorial Sloan Kettering Cancer Center and followed the National Institutes of Health guidelines for animal welfare.

### Subcutaneous tumor implantation and TMZ plus TAL treatment

5 × 10^6^ cells/mouse of MDA-MB-468 cells or 1.5 × 10^6^ cells/mouse PDX WHIM25 were suspended in 100 μL of 1:1 PBS/matrigel basement membrane matrix (Corning) and injected subcutaneously on the flank of female NSG mice (NOD.Cg-Prkdcscid Il2rgtm1Wjl/SzJ; The Jackson Laboratory) that were 6 to 8 weeks old (n=5). Tumor volumes were calculated from manual caliper measurements as volume = π/6 (length × width × width). Once tumor volumes reached approximately 150 mm^3^, mice were treated via oral gavage with vehicles, or in combination (TAL 0.2mg/kg daily and TMZ 6 mg/kg every 4 days).

### Orthotopic tumor implantation and TMZ plus TAL treatment

5 × 10^6^ cells/mouse of MCF7 cells expressing wtp53, MDA-MB-468 cells expressing mtp53 R273H, mtp53 R273H(Δ347-393), or mtp53 R273Hfs387 were implanted into the mammary fat pad of female NSG mice (The Jackson Laboratory) that were 6 to 8 weeks old (n=5). Tumor volumes were calculated from manual caliper measurements as volume = π/6 (length × width × width). For TMZ plus TAL treatment, once tumor volumes reached approximately 150 mm^3^, mice were treated via oral gavage with vehicles, or in combination (TAL 0.2mg/kg daily and TMZ 6 mg/kg every 4 days). For MCF7 study, animals were provided with drinking water with containing 8 μg/ml 17β-estradiol (Millipore Sigma).

### Circulating tumor cell analysis

CTCs were isolated as described previously [23, 52]. At the endpoint of the experiment, animals were euthanized and blood samples were collected by cardiac puncture and stored in microcentrifuge tubes coated with sodium heparin (Cat# S1346, Selleckchem) prior to CTC isolation. Whole blood was subjected to centrifugation and the buffy coat layer was then collected and subjected to red blood cell (RBC) lysis to remove residual RBCs. Cells were washed with PBS and stained with phycoerythrin (PE) conjugated mouse anti-human EGF Receptor antibody (Cat#555997; BD Pharmigen). Stained cells were fixed in 1% paraformaldehyde in PBS. Flow cytometric analysis was performed using a FACScan device (BD Biosciences), and event counting was gated based on cell size and EGFR intensity. The number of CTCs was obtained by dividing the number of positive events by individual blood volume. Statistical analysis was generated using Prism 9.0 software (GraphPad). In the box-and-whisker plots, each dot represents one mouse.

### Tissue processing and quantification of lung metastases

Animal tissues were harvested, fixed in 10% buffered formalin, and embedded in paraffin. Sections of lungs were cut at 5 μm and immunohistochemistry (IHC) staining of human-specific mitochondrial antigen (Hu-MITO) was performed. Stained slides were scanned on an Olympus VS200 slide scanner and digital images were acquired in a. VSI format. Using these scanned slides, quantitative analysis of Hu-MITO immunostaining was performed using QuPath software version 0.7.0 (https://qupath.github.io/) [53]. A region of interest (ROI) was manually delineated around each lung section on each slide, and positive cell detection was applied. The best Qupath positive cell detection settings were determined by a pathologist at LCP (MIA) and these settings were used for all slides. Annotation measurements, expressed as the percentage of positive cells (detected positive cells/detected negative cells), were subsequently exported and tabulated in an Excel spreadsheet.

### Tumor protein extraction

The animals were euthanized by CO_2_(g) asphyxiation and tumors were removed and snap frozen. Protein extraction from frozen tumor was described previously [3]. Tumor chromatin extraction was carried out by Chromatin Extraction Kit from Abcam (Cat# ab117152).

### DNA fiber combing with S1 nuclease

DNA combing with S1 nuclease was performed as previously described with modifications [54, 55]. Exponentially growing cells pulse-labeled with 100 μM CldU (5-chloro-29-deoxyuridine, Cat# C6891; Sigma) in cell culture media for 30 min and washed three times with PBS. Cells then labeled with 100 μM IdU (5-Iodo-29-deoxyuridine, Cat# I7125; Sigma) for 60 min with or without 10 μM PARGi (Cat# HY-133531; MCE). Cells were then collected by trypsinization and 5 × 10^5^ cells in 45 ul trypsin was prewarmed to 50°C for 10 seconds and embedded into 45 ul agarose plug (Cat#IB70050; IBI). The plugs were incubated overnight at 42°C in proteinase K lysis buffer (2 mg/ml proteinase K, 10 mM Tris-HCl pH 7.5, 100 mM EDTA, 0.5% SDS, and 20 mM NaCl). DNA plugs were washed two times for 1 h in TE50 buffer (10 mM Tris-HCl pH 7.5, 50 mM EDTA, and 100 mM NaCl) followed by two times for 1 h in TE buffer with 100 mM NaCl (10 mM Tris-HCl pH 7.5, 1 mM EDTA, and 100 mM NaCl). The plugs were melted at 68°C for 20 min in 1 ml 0.5M MES pH 5.5 (Cat# 475893; Millipore) and cooled to 42°C for 10 min. The samples were incubated at 42°C overnight with the addition of 3 μl of β-agarose (Cat# M0392L; NEB) dissolved in 100 µl MES solution. The DNA mix was then gently poured into combing reservoirs containing 1.2 ml MES supplemented with 2 mM Zn(O2CCH3)2 and either S1 nuclease for final concentration of 40 U/ml (Cat# EN0321; Thermo Fisher) or S1 nuclease dilution buffer and incubated for 30 min at room temperature. The samples were stored at 4°C for two days and genomic DNA was combed onto salinized coverslips (Cat# Po-104 90 802; AutoMate Scientific) using the combing machine from Genomic Vision and baked for 2 h at 68°C. For immunostaining, DNA was denatured with fresh 0.5 M NaOH /1 M NaCl solution for 8 min at RT. Coverslips were then washed 2 min with PBS for three times, dehydrated with 70% and 100% ethanol for 5 min each, and dry for 30 min at room temperature. Coverslips were blocked with 10% goat serum in PBS at 37°C for 30 min. DNA fibers were immuno-stained with rat anti-BrdU for CldU detection (1:45, Cat# Ab6326, Abcam) and mouse-anti-BrdU for IdU detection (1:10, Cat# 347580, BD Biosciences) in 5% BSA in PBS for 1 h at 37°C, washed 5 min two times with PBS-0.01% Tween-20, and 1 time PBS. Coverslips were incubated with goat anti-rat Alexa Fluor 488 (1:100, Cat# A11006, Thermo Fisher) and goat anti-mouse Alexa Fluor 594 (1:100 Cat# A11005; Thermo Fisher) for 60 min at 37°C. Coverslips were washed 5 min two times with PBS-0.01% Tween-20, and 1 time PBS. Coverslips were dried before mounting on microscope slide with Prolong Diamond Antifade (Cat# P36965; Thermo Fisher). Images were taken using Nikon A1 confocal microscope with 60x objective oil immersion, acquired by Nikon elements. IDU track lengths were measured by ImageJ software.

### Differential Gene Expression (Bulk RNA-Seq)

EdgeR/DeSeq2 for differential gene expression analysis was performed. TMZ + TAL as the treatment group compared to the control group. Multiple test correction was done with Benjamin-Hochberg and an adjusted p-value of 0.05 was applied to identify significantly expressed genes. Pathway analysis was performed using Gene Set Enrichment Analysis (GSEA) on genes ranked by signed log2 fold change. Enrichment was evaluated against the MSigDB Hallmark gene sets. Pathways with an FDR q-value below the 0.05 threshold were considered significantly enriched. Enrichment direction (up- or down-regulated) was determined by the normalized enrichment score (NES).

## Supporting information

Supplementary RNA Sequence Data

## Acknowledgments and funding sources

We thank Anna Boers, Yasmin Hussein and members of the Bargonetti team for their efforts in support of this research. We would like to thank Allen Annis former SVP of Aileron Therapeutics for providing us with ALRN. We also thank Jacqueline Candelier of The Laboratory of Comparative Pathology at Memorial Sloan Kettering Cancer Center and Joon Kim from Flow Cytometry Facility at Hunter College for circulating tumor cells detection. We thank Zhengming Chen for expert assistance with biostatistical analysis.This work was supported by The Breast Cancer Research Foundation BCRF-20-011, BCRF-21-011, BCRF-22-011, BCRF-23-011, and BCRF-24-011 to J. Bargonetti, National Cancer Institute of the National Institutes of Health under Award Number R01CA239603 to J. Bargonetti, and by the TUFCCC/HC Regional Comprehensive Cancer Health Disparity Partnership from the NIH U54 CA221704 and V. Chavez was supported by RISE program at Hunter College funded by NIH grant R25GM060665 to Hunter College.

