## Supplemental Figures 1 to 5 for "Mutant p53 Directs PARP to Regulate Replication Stress and Drive Breast Cancer Metastasis"

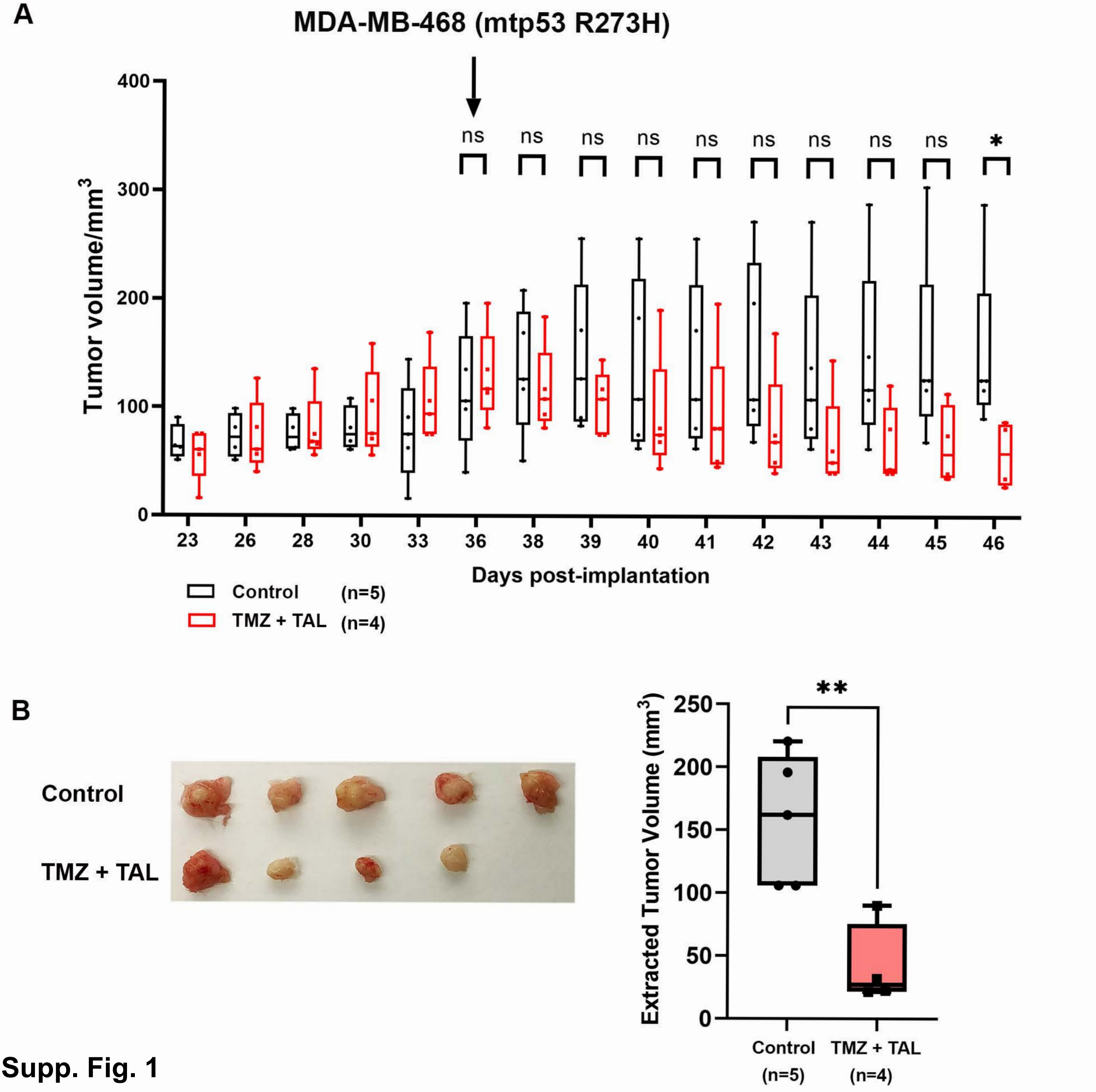
A
MDA-MB-468 (mtp53 R273H)
ns
*
Tumor volume/mm3
400
300
200
100
0
23
26
28
30
33
36
38
39
40
41
42
43
44
45
46
Days post-implantation
Control (n=5)
TMZ + TAL (n=4)
B
Control
TMZ + TAL
**
Extracted Tumor Volume (mm3)
250
200
150
100
50
0
Control (n=5)
TMZ + TAL (n=4)

Supp. Fig. 1

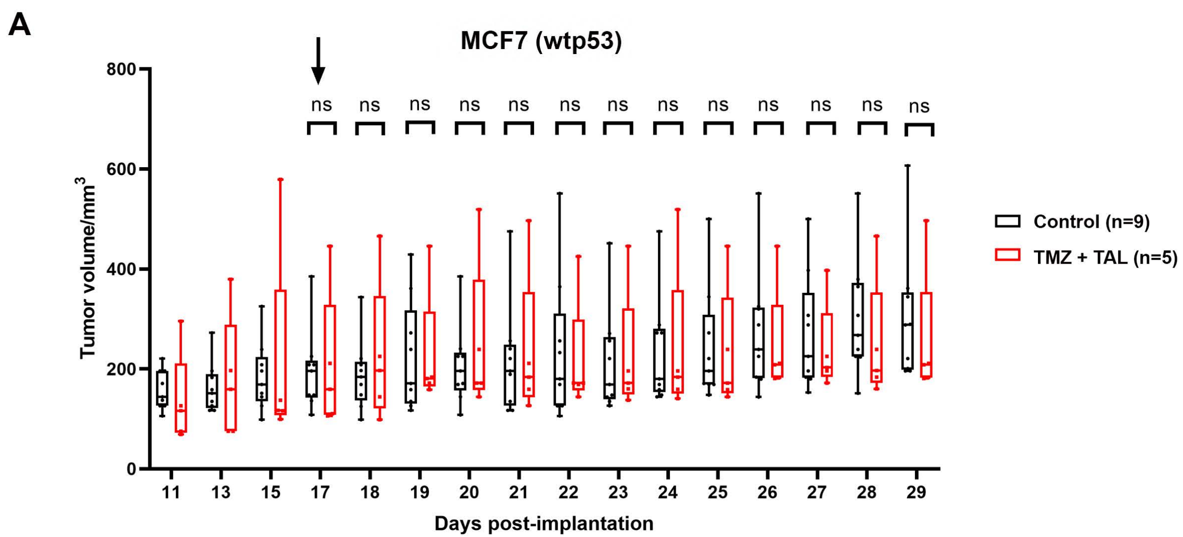
A
MCF7 (wtp53)
ns
Tumor volume/mm3
800
600
400
200
0
11 13 15 17 18 19 20 21 22 23 24 25 26 27 28 29
Days post-implantation
Control (n=9)
TMZ + TAL (n=5)

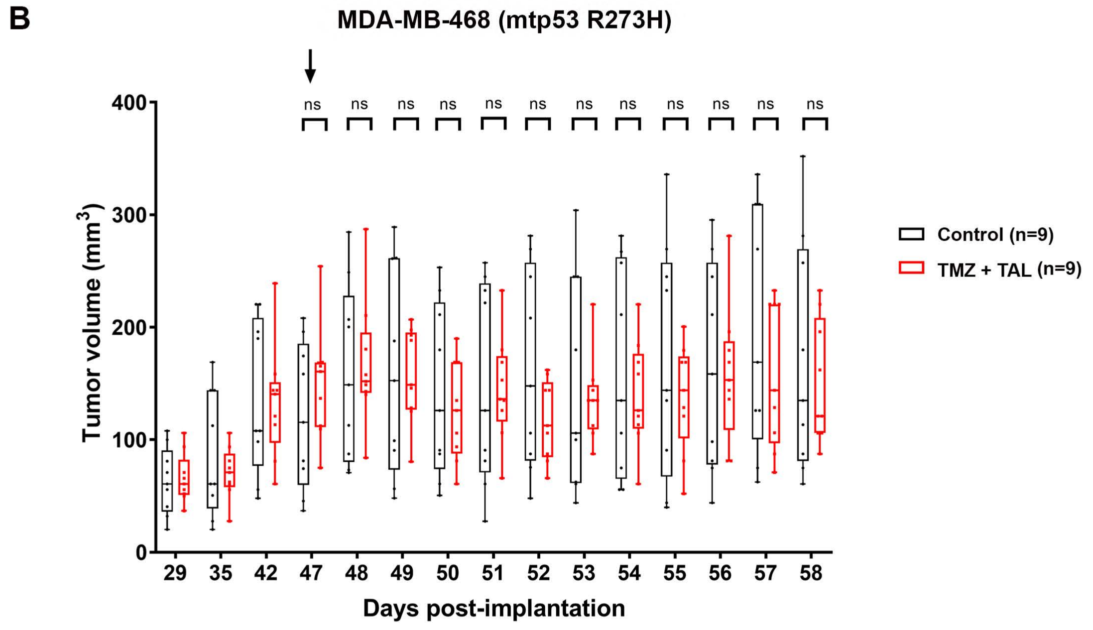
B
MDA-MB-468 (mtp53 R273H)
ns
Tumor volume (mm3)
400
300
200
100
0
29 35 42 47 48 49 50 51 52 53 54 55 56 57 58
Days post-implantation
Control (n=9)
TMZ + TAL (n=9)

Supp. Fig. 2

**A**

WHIM25 (mtp53 R273H)

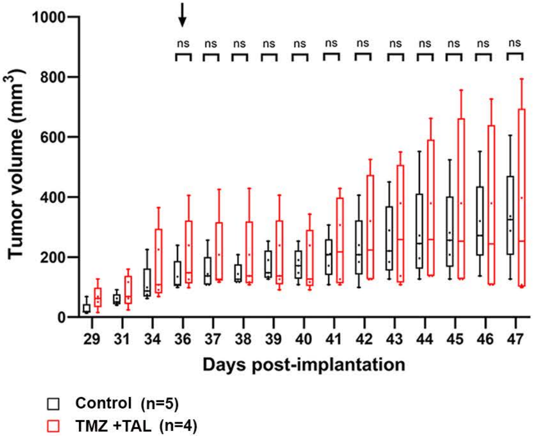


WHIM6 (Loss of p53 expression)

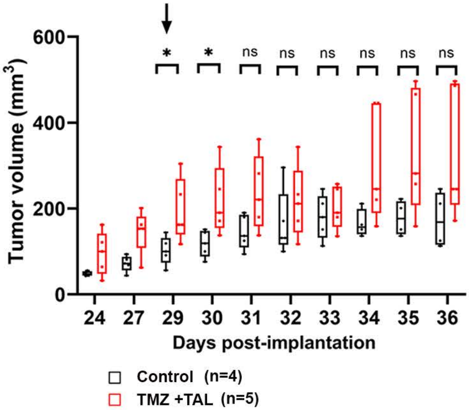


**B**

WHIM25 (mtp53 R273H)

Control

TMZ +TAL

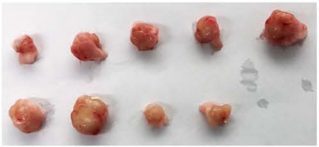

WHIM6 (Loss of p53 expression)

Control

TMZ +TAL

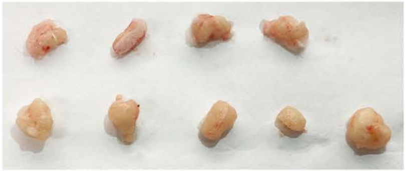

**Supp. Fig. 3**

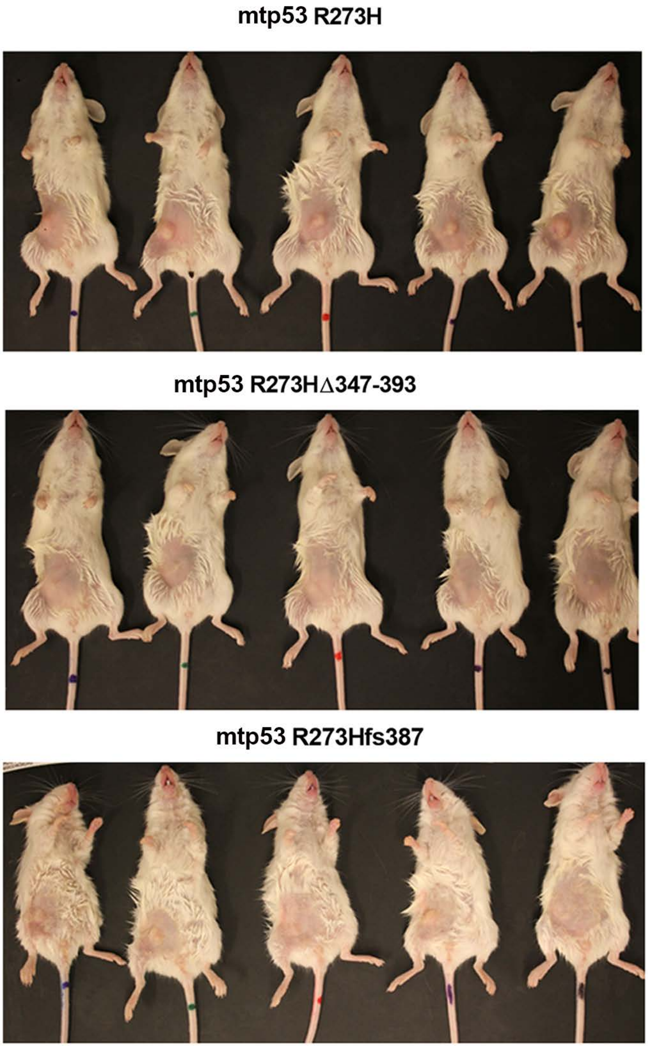


Supp. Fig. 4

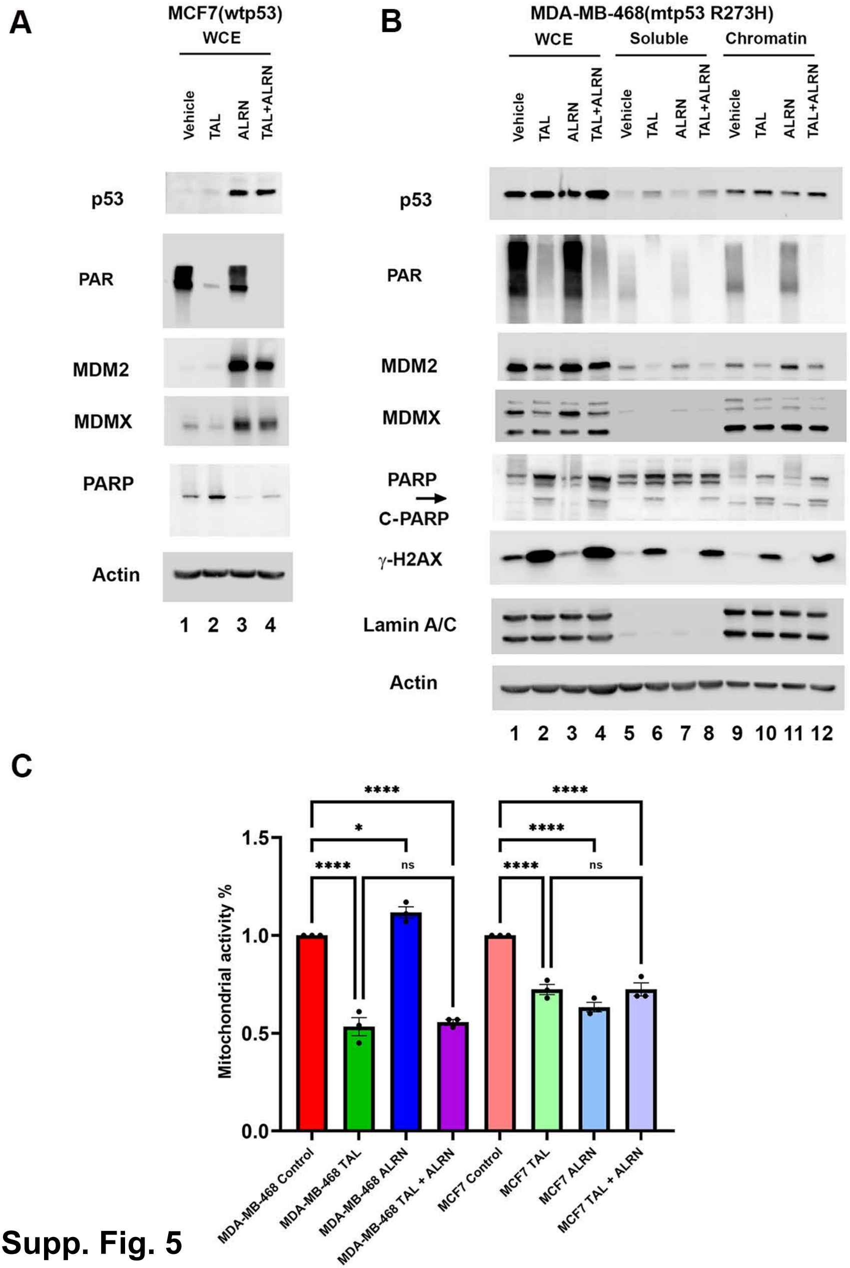
A
MCF7(wtp53)
WCE
Vehicle
TAL
ALRN
TAL+ALRN
p53
PAR
MDM2
MDMX
PARP
Actin
1 2 3 4
B
MDA-MB-468(mtp53 R273H)
WCE
Soluble
Chromatin
Vehicle
TAL
ALRN
TAL+ALRN
p53
PAR
MDM2
MDMX
PARP
C-PARP
γ-H2AX
Lamin A/C
Actin
1 2 3 4 5 6 7 8 9 10 11 12
C
Mitochondrial activity %
1.5
1.0
0.5
0.0
****
*
****
ns
MDA-MB-468 Control
MDA-MB-468 TAL
MDA-MB-468 ALRN
MDA-MB-468 TAL + ALRN
MCF7 Control
MCF7 TAL
MCF7 ALRN
MCF7 TAL + ALRN

Supp. Fig. 5

**D**

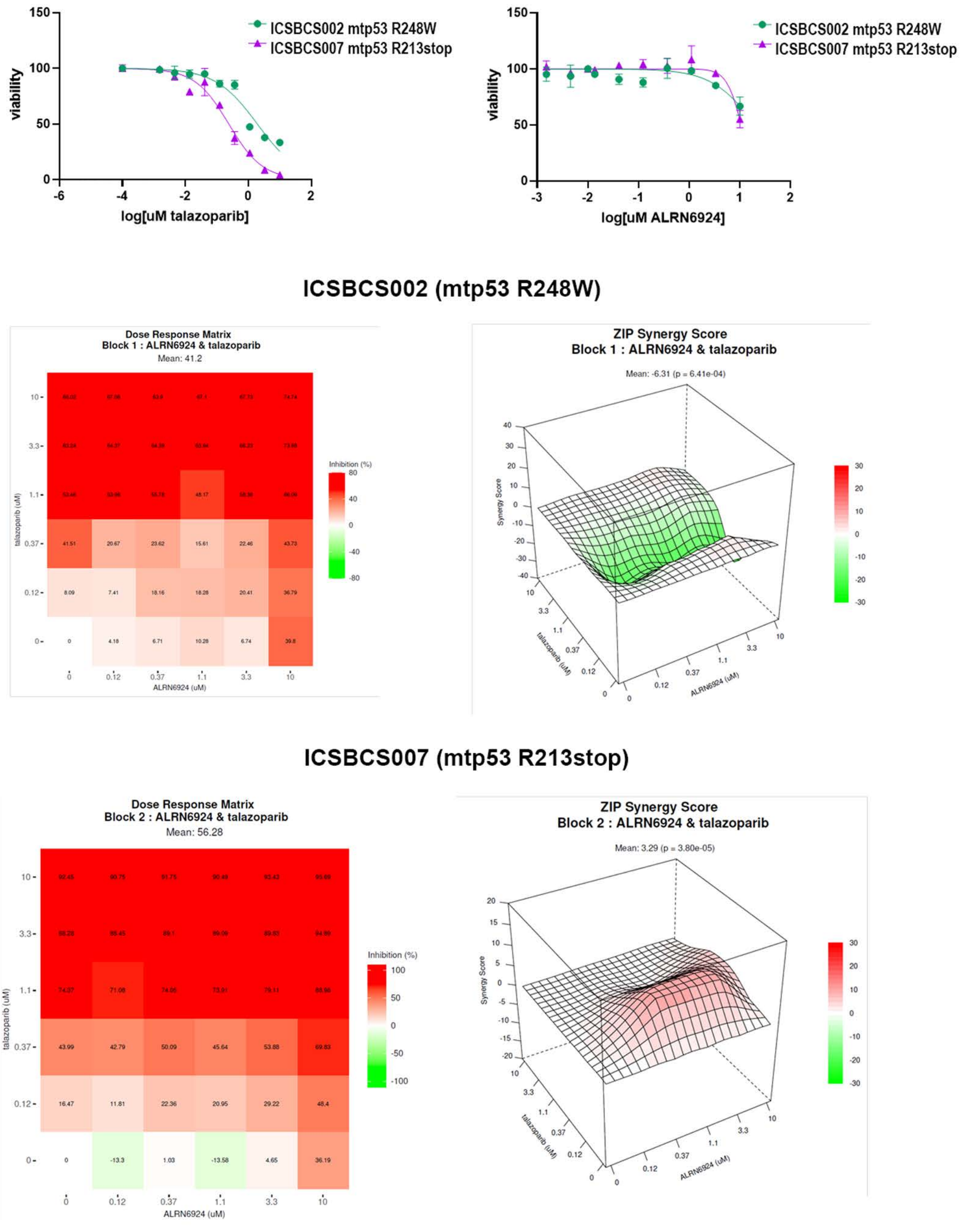
ICSBCS002 mtp53 R248W
ICSBCS007 mtp53 R213stop
viability
log[uM talazoparib]
log[uM ALRN6924]
ICSBCS002 (mtp53 R248W)
Dose Response Matrix
Block 1 : ALRN6924 & talazoparib
Mean: 41.2
Inhibition (%)
talazoparib (uM)
ALRN6924 (uM)
ZIP Synergy Score
Block 1 : ALRN6924 & talazoparib
Mean: -6.31 (p = 6.41e-04)
Synergy Score
ICSBCS007 (mtp53 R213stop)
Dose Response Matrix
Block 2 : ALRN6924 & talazoparib
Mean: 56.28
Inhibition (%)
talazoparib (uM)
ALRN6924 (uM)
ZIP Synergy Score
Block 2 : ALRN6924 & talazoparib
Mean: 3.29 (p = 3.80e-05)
Synergy Score

**Supp. Fig. 5**
