## Supplementary Figures Legend for "Mutant p53 Directs PARP to Regulate Replication Stress and Drive Breast Cancer Metastasis"

**Supplementary figure legends**

**Supplementary Figure 1 Daily measurement of tumor volume and endpoint tumor size of subcutaneous MDA-MB-468 tumors with TMZ plus TAL treatment**

(A) MDA-MB-468 cells were subcutaneous implanted in the flank of female NSG mice. Once tumor volumes reached approximately 150 mm3, mice were randomized to treatment via oral gavage either with vehicles (Control), or in combination of talazoparib 0.2mg/kg daily and temozolomide 6 mg/kg every 4 days (TMZ + TAL) treatment group. Primary tumor volumes of MDA-MB-468 vehicle (Control) and MDA-MB-468 (TMZ +TAL) mice were measured using calipers as volume = π/6 (length × width × width). Arrow indicated the first day of treatment. (B) Picture of Endpoint primary tumors from Control and TMZ + TAL group and quantification of endpoint tumor volumes. In the box-and-whisker plots, each dot represents one mouse, median values are represented by horizontal lines. ns., not significant, *P < 0.05, **P < 0.01, Multiple Mann-Whitney test.

**Supplementary Figure 2 Daily measurement of orthotopic tumor volume of MCF7 and MDA-MB-468 tumors with TMZ plus TAL treatment**

(A) MCF7 and (B) MDA-MB-468 cells were orthotopic implanted into the mammary fat pad of female NSG mice. Once tumor volumes reached approximately 150 mm3, mice were randomized to treatment via oral gavage either with vehicles (Control), or in combination of talazoparib 0.2mg/kg daily and temozolomide 6 mg/kg every 4 days (TMZ + TAL) treatment group. Primary tumor volumes of MDA-MB-468 vehicle (Control) and MDA-MB-468 (TMZ + TAL) mice were measured using calipers as volume = π/6 (length × width × width). Arrow indicated the first day of treatment. In the box-and-whisker plots, each dot represents one mouse, median values are represented by horizontal lines. ns., not significant, Multiple Mann-Whitney test.

**Supplementary Figure 3 Daily measurement of tumor volume of subcutaneous of PDX WHIM25 and WHIM6 with TMZ plus TAL treatment**

(A) PDX WHIM25 or WHIM6 cells were subcutaneous implanted in the flank of female NSG mice. Once tumor volumes reached approximately 150 mm3, mice were randomized to treatment via oral gavage either with vehicles (Control), or in combination of talazoparib 0.2mg/kg daily and temozolomide 6 mg/kg every 4 days (TMZ + TAL) treatment group. Primary tumor volumes vehicle (Control) and TMZ +TAL mice were measured using calipers as volume = π/6 (length × width × width). Arrow indicated the first day of treatment. In the box-and-whisker plots, each dot represents one mouse, median values are represented by horizontal lines. ns., not significant, Multiple Mann-Whitney test. (B) Picture of endpoint primary tumors from Control and TMZ + TAL group.

**Supplementary Figure 4 Endpoint tumor picture of orthotopic MDA-MB-468 tumors**

MDA-MB-468 mtp53, mtp53 R273H(Δ347-393) and mtp53 R273Hfs387 cells were orthotopic implanted into the mammary fat pad of female NSG mice. At the endpoint, the tumor development was imaged.

**Supplementary Figure 5 Combination treatment with ALRN-6924 and PARP inhibitor talazoparib in breast cancer cells and patient-derived tumor organoids PDTOs**

Whole cell lysate (WCE) were prepared from MCF7 (A) and whole cell lysate (WCE), soluble and chromatin fractions were prepared from MDA-MB-468 (B) treated with either vehicle (DMSO), 10 μM talazoparib (TAL), 10 μM ALRN-6924 (ALRN), or combination (TAL + ALRN) for 24 h. Protein levels of p53, PAR, MDM2, MDMX and PARP were determined by Western blot analysis. (C) MTT assay was performed to measure mitochondrial activity. MDA-MB-468 or MCF7 cells were seeded in 12-well plates and attached overnight. Cells were treated with either 10 μM talazoparib or 10 μM ALRN-6924, or both for 48 hours. The absorbance was quantified by measuring the absorbance at 550 nm subtracted from the absorbance at 620 nm. All MTT data are represented as mitochondrial dehydrogenase activity as percentage of a dimethyl sulfoxide (DMSO) vehicle treated control. ns., not significant *P < 0.05, **P < 0.01, ***P <0.001, ****P < 0.0001, two-way ANOVA test followed by Tukey’s post-hoc test. (D) Talazoparib or ALRN-6924 response curves for breast cancer organoids PDTO mtp53 R248W (ICSBCS002) and mtp53 R213stop (ICSBCS007). Cells were plated in 384 well plates and inhibitors were added after 3 days. After 4-day incubation viability was determined using Cell Titer Glo 3D reagent. Synergy maps for talazoparib-ALRN were calculated using the SynergyFinder web application with the ZIP synergy model (red indicates a synergistic effect, white an additive effect, and green an antagonistic effect).
